# The impact of Library Size and Scale of Testing on Virtual Screening

**DOI:** 10.1101/2024.07.08.602536

**Authors:** Fangyu Liu, Olivier Mailhot, Isabella S. Glenn, Seth F. Vigneron, Violla Bassim, Xinyu Xu, Karla Fonseca-Valencia, Matthew S. Smith, Dmytro S. Radchenko, James S. Fraser, Yurii S. Moroz, John J. Irwin, Brian K. Shoichet

**Affiliations:** Dept. of Pharmaceutical Chemistry, University of California, San Francisco, San Francisco CA 94143, USA; Dept. of Bioengineering and Therapeutic Sciences, University of California, San Francisco, San Francisco CA 94143, USA; Enamine Ltd., Kyiv, 02094, Ukraine; Chemspace (www.chem-space.com), Chervonotkatska Street 85, Kyїv 02094, Ukraine; Taras Shevchenko National University of Kyїv, Volodymyrska Street 60, Kyїv 01601, Ukraine

## Abstract

Virtual libraries for ligand discovery have recently increased 10,000-fold, and this is thought to have improved hit rates and potencies from library docking. This idea has not, however, been experimentally tested in direct comparisons of larger-vs-smaller libraries. Meanwhile, though libraries have exploded, the scale of experimental testing has little changed, with often only dozens of high-ranked molecules investigated, making interpretation of hit rates and affinities uncertain. Accordingly, we docked a 1.7 billion molecule virtual library against the model enzyme AmpC β-lactamase, testing 1,521 new molecules and comparing the results to the same screen with a library of 99 million molecules, where only 44 molecules were tested. Encouragingly, the larger screen outperformed the smaller one: hit rates improved by two-fold, more new scaffolds were discovered, and potency improved. Overall, 50-fold more inhibitors were found, supporting the idea that there are many more compounds to be discovered than are being tested. With so many compounds evaluated, we could ask how the results vary with number tested, sampling smaller sets at random from the 1521. Hit rates and affinities were highly variable when we only sampled dozens of molecules, and it was only when we included several hundred molecules that results converged. As docking scores improved, so too did the likelihood of a molecule binding; hit rates improved steadily with docking score, as did affinities. This also appeared true on re-analysis of large-scale results against the σ2 and dopamine D4 receptors. It may be that as the scale of both the virtual libraries and their testing grows, not only are better ligands found but so too does our ability to rank them.

## Introduction

With the advent of ultra-large, make-on-demand (“tangible”) libraries, available chemical space has increased from about 3.5 million to over 38 billion (https://enamine.net/compound-collections/real-compounds). While the size of the new libraries can seem daunting, recent studies suggest that structure-based docking prioritizes potent ligands from within it, with affinities often in the mid-nanomolar and sometimes high picomolar range^1–11^. Docking the new libraries seems to improve hit rates, affinities, and chemotype novelty versus smaller libraries^12,13^, suggesting that bigger libraries are better for virtual screening. This is supported by simulations that show that as libraries grow, the best molecules fit ever better to protein binding sites^14^. Exactly how large libraries affect these key outcomes versus smaller libraries remains to be tested experimentally in side-by-side studies.

Further clouding the issue is the scale of experimental testing of molecules prioritized from virtual screens. Irrespective of whether million-scale or billion-scale libraries are virtually screened, rarely are more than several dozen molecules synthesized and tested experimentally^3,6–8^. While these are high-ranking, they are picked from among a much larger pool of similarly ranked molecules. From the hit rates of these screens (number active/number-tested), it has been inferred that there are likely hundreds-of-thousands or even millions of potential ligands in the libraries that remain untested, but this has not been probed experimentally^1^. As important, the few molecules tested make the results subject to the statistics of small numbers. Said another way, it is not clear that we can have full confidence in hit rates, affinity ranges, and the likelihood of discovering new chemotypes—all key docking outcomes—when testing only a few dozen compounds.

Here we begin to investigate these questions quantitatively. **First,** to explore the impact of library size on docking outcome, we screened over 1.7 billion molecules for inhibitors of the model enzyme AmpC β-lactamase^1,15–20^ and compared the results to a previous screen on the same enzyme using essentially the same method where only 99 million molecules were docked^1^. These smaller and larger screens were compared by hit rates, inhibitor affinities, and the number of novel chemotypes discovered. **Second**, we synthesized and tested 1,521 compounds for AmpC inhibition, rather than the 44 originally tested in the smaller library campaign^1^, and asked if the number of inhibitors found scaled with number of top-ranking molecules investigated, something that has until now simply been an implication of large library docking. **Third**, with these observations in hand, we examined the sensitivity of docking hit rates and hit affinities to the scale of experimental testing by sub-sampling smaller sets from the larger one; this has implications for how we should understand docking hit rates and affinities, and how we should scale these experiments in the future.

**Fourth,** we investigate how hit rate is predicted by docking score, and whether we might expect better molecules to be found as libraries continue to expand into the tens of billions of molecules and beyond^5,21^. **Finally,** the scale of the experimental testing here allows us to investigate potential correlations between docking rank and affinity category (high, mediocre, poor). We will argue that the answers emerging from this large-scale study support further expansion of docking libraries into the trillions of compounds range, and, perhaps surprisingly, a re-investment in docking scoring functions to optimize what is now a loose correlation between docking rank and affinity category.

## Results

### Selection, synthesis, and testing of 1521 docking hits for AmpC

With a quantitative spectrophotometric assay, ability to determine inhibitor-protein crystal structures, and status as a primary antibiotic resistance mechanism, AmpC β-lactamase has lent itself to multiple structure-based and high-throughput screening campaigns for inhibitor discovery^15–20^, including with ultra-large libraries^1^, making it a good system to test the impact of library size on virtual screening. In a previous docking screen of 99 million molecules against the enzyme, 44 high-ranking molecules, topologically unrelated to previously known scaffolds, were prioritized for synthesis and testing. This revealed five new inhibitors with affinities ranging from 1.3 µM to 400 µM, a hit rate of 11% using this range of activity^1^. Using essentially the same docking method, here we screened a 1.7 billion molecule library against the same AmpC active site. Molecules from across the docking scoring range, 838,672,414 in total ranking from -117.35 kcal/mol (best scores) to -28 kcal/mol (worst scores), were considered as candidates for experimental testing. These were organized into bins of resolution ranging from 2 to 4 kcal/mol among the lower (better) scores to 8 kcal/mol among the higher (worse) scores. Up to 25,000 molecules were selected per bin, by rank order (for the lower and better energy bins, this amounted to all the molecules in the bin). Molecules topologically similar to known inhibitors, with ECFP4-based Tc > 0.5, were excluded, as were those with more than one unsatisfied hydrogen bond donor and more than six hydrogen bond acceptors—such molecules exploit known gaps in the DOCK3.8 scoring function^22^. The remaining 193,878 molecules were clustered by *Tc* = 0.32 based on the interaction fingerprinting^23^, resulting in 80,767 cluster heads. In previous simulations^14^ and experiments^4^, we had found molecules with artifactually favorable score concentrated among the top-ranking docked molecules. Here too, we observed molecules that achieved scores much higher than one would expect from the overall distribution; this problem became more acute as the library grew (**Extended Data** Fig. 1). We chose to ignore these molecules for experimental testing. The origins of these molecules, and their experimental confirmation as docking artifacts, is explored in a separate study [Wu, 2024].

Overall, 2,089 cluster-heads, all topologically dissimilar to one another and to known inhibitors, were chosen for synthesis and testing. Of these, 1,521 were successfully synthesized (a fulfillment rate of 73%). Manual inspection (“manual-picked”) from among the better scoring bins (-100.58 to -79 kcal/mol) accounted for 687 of these compounds, and another 1,292 molecules were chosen based on rank alone (“auto-picked”), with 458 molecules occurring in both sets.

All molecules were initially tested at 200, 100, and 40 µM concentrations for AmpC inhibition^1,16,20^. Of the 1,447 experimentally well-behaved molecules, 1,296 were among the top scoring 1% of the docked molecules, the same cut-off used in the 99 million molecule screen (the rest were spread out among lower ranks and were selected to test hit rate versus score dependence). Of these 1,296 compounds, 171 had an apparent Ki < 166 µM, based on the three-point inhibition numbers and assuming competitive inhibition (see below), while another 124 had apparent Ki values between 166 and 400 µM. Concentration-response curves were measured for 17 compounds across this potency range. The IC50 values from these full curves corresponded well to those predicted by the three-point inhibition numbers (**Extended Data Table 1, Extended Data** Fig. 2). For seven of the new inhibitors, each in a different chemotype family, we determined full Ki values and mechanism by Lineweaver-Burk analysis (**Extended Data** Fig. 3). All seven were competitive inhibitors, consistent with docking to the AmpC active site, with Ki values ranging from 0.7 to 4.6 µM (**Extended Data** Fig. 3). Accordingly, we modeled all of the new inhibitors as competitive, consistent with the x-ray crystal structures determined for five of the new inhibitors, which were all observed to bind in the β-lactamase active site (**Fig. 1**). With this assumption, Ki values ranged from 464 to 0.46 µM^24^, with substantial representation across this range of affinities (**Fig. 2**). All assays included 0.01% Triton X-100, diminishing the likelihood of artifact from colloidal aggregation^18,25^. For further confidence, 140 of the inhibitors were checked for particle formation by dynamic light scattering (DLS) ^25–27^; no signs of colloid-like particle formation were detected among any of them at relevant concentrations (**Extended Data Table 2**).

**Fig. 1.**
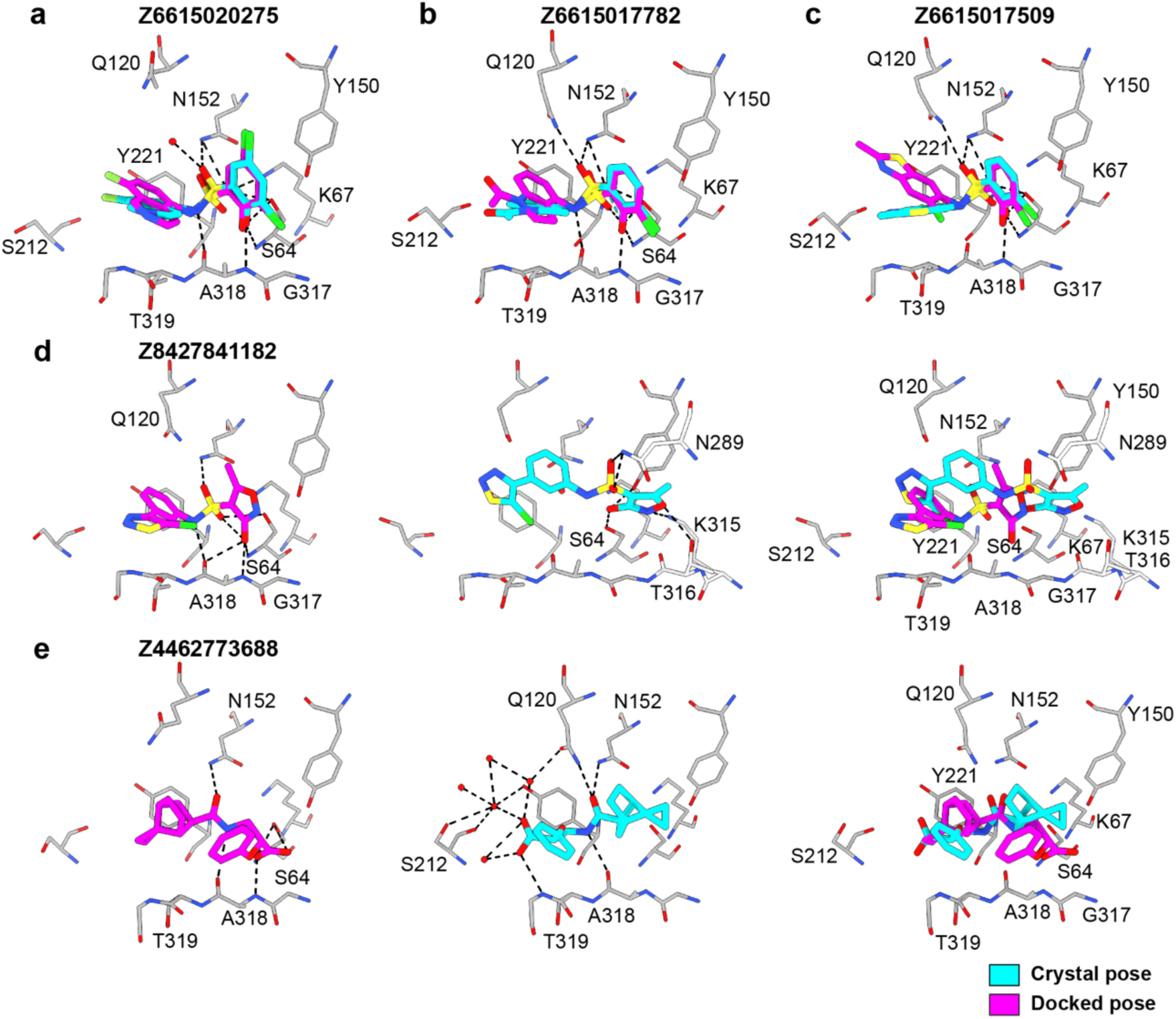
Superposition of the crystallographic and docking poses of the new AmpC inhibitors. Crystal structures (carbons in cyan) and docked poses (carbons in magenta) of the inhibitors. AmpC carbon atoms are in grey, oxygens in red, nitrogens in blue, sulfurs in yellow, chlorides in green, and fluorides in light blue. Hydrogen bonds are shown as black dashed lines. **a-c,** AmpC in complex with **Z6615020275** (r.m.s.d to crystal structure 0.79 Å, Ki 2 uM), **Z6615017782** (r.m.s.d = 0.97 Å, 1.5 uM) and **Z6615017509** (r.m.s.d = 3.14 Å, 0.86 nM). The overlay of the crystal and docked poses are shown. **d-e,** AmpC in complex with **Z8427841182** (r.m.s.d = 4.73 Å, 36 uM) and **Z4462773688** (r.m.s.d = 5.61 Å, 325 uM). The docked poses (left panel), crystal poses (middle panel) and the overlay of the docked and crystal poses are shown (right panel).

**Fig. 2.**
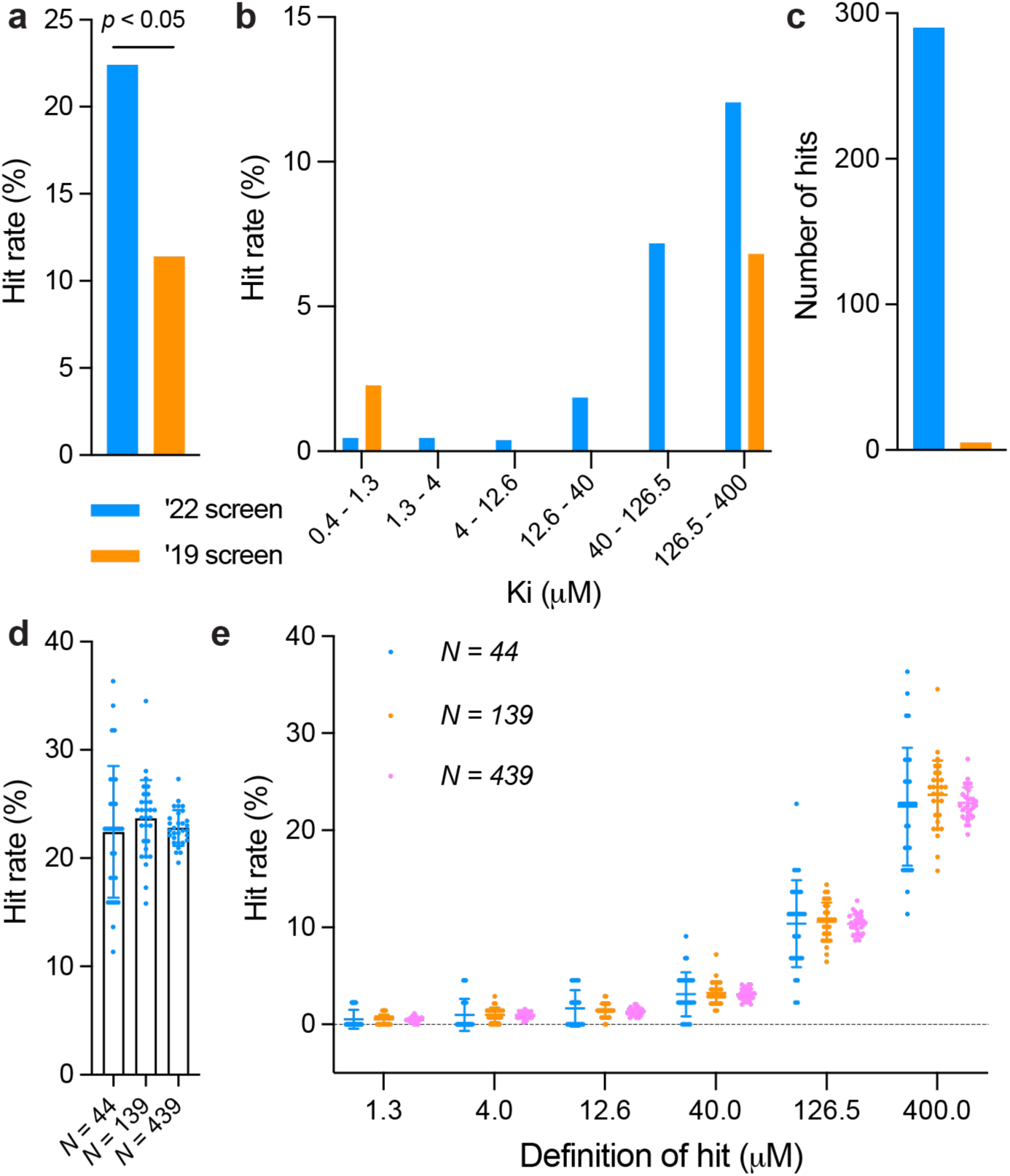
Larger-scale docking and testing increases hit rates and reduces uncertainty. **a,** The hit rates (number of actives/total tested) of the **1.7 B**illion screen (blue bar) versus the **99 M**illion screen (orange bar). **b,** Hit rates by different affinity bins in ‘22 screen and ‘19 screen. **c,** Number of hits (number of actives) of the **1.7 B** screen (blue bar) versus the **99 M** screen (orange bar). **d,** The impact of randomly purchasing 44, 139, 439 molecules out of 1,296 molecules for testing on hit rates. Each sample size is randomly drawn 30 times and the resulting hit rates were plotted. The error bars represent SDs of the hit rates. The hit rates are 22.42 ± 6.08% (*N* = 44), 23.67 ± 3.54% (*N* = 139) and 22.80 ± 1.65% (*N* = 439). **e,** The impact of randomly purchasing 44, 139, 439 molecules out of 1,296 molecules for testing on hit rates with different affinity cutoffs. Each sample size is drawn 30 times and the resulting hit rates were plotted. The error bars represent SDs of the hit rates.

### Docked versus crystallographic geometries

To investigate how docking predicted geometries corresponded to experimentally-determined ligand poses, the structures of five of the new inhibitors were determined by x-ray crystallography, with resolutions varying from 1.6 to 2.9 Å (**Extended Data Table 3**). Unambiguous electron density allowed us to confidently model the positions of the new inhibitors in the enzymes’ active site. For **Z6615020275** (1.3 µM; **Fig. 1a**), **Z6615017782** (0.95 µM; **Fig. 1b**) and **Z6615017509** (0.86 µM; **Fig. 1c**), the docked and experimental structures superimposed with a 0.79, 0.97, and 3.14 Å root mean square deviation (RMSD) respectively, with the differences in position stemming from deviations of non-warhead groups binding distally in the site. For two weaker inhibitors, the crystal structures had larger deviations from the docking predictions. While the crystallographic pose of **Z8427841182** (36 µM) hydrogen-bonded with many of the same residues predicted in the docking pose, the crystallographic geometry was shifted in the site and the RMSD was high at 4.73 Å RMSD. This hydroxy-isoxazole, a close analog of the original 36 µM docking hit **Z6615146331** that had resisted facile crystallization (**Extended Data Table 1**), represents a previously unknown warhead for AmpC. Meanwhile, the crystallographic pose of the 323 µM **Z4462773688**, an unprecedented bicyclo-alkyl carboxylate, bound in a geometry flipped from that anticipated by docking, leading to an RMSD of 5.61 Å (**Fig. 1e**). **‘6631** and **‘3688** are two examples of the 44 inhibitors found in this campaign that sample not only novel topologies, but also novel warheads for AmpC.

### Hit rates are higher from the larger vs the smaller library screens

The overall hit rate (number experimentally active/number experimentally tested) from the 1.7 billion molecule docking screen was 22.4% (290 actives/1,296 high-ranking tested). This hit rate is significantly higher than that from 99 million molecule docking screen, which was 11.4% (p-value < 0.05) (**Fig. 2a**). The hit rate difference stands out even more strongly when considered across affinity ranges. Most of the actives from the 99 million molecule screen had apparent Ki values between 126.5 and 400 µM (**Fig. 2b**), with one inhibitor found in the 1 to 10 µM range, and none found in the intermediate ranges. Conversely, from the 1.7 billion library each half-log affinity bin is well-populated by active molecules. The higher hit rate from the larger library is consistent with the idea that as the virtual libraries grow, ever more plausible molecules are fortuitously sampled and prioritized by molecular docking.

### Hit rate variability and ligand affinity ranges

While hit rate is a fair way to compare the two screens, naturally, the raw number of hits was far greater from the larger library (290 active from 1.7 billion screened versus 5 actives from 99 million screened, **Fig. 2c**), where 29-fold more high-ranking molecules were tested. Qualitatively this explains why all half-log affinity bins were well-populated from the larger library, whereas this was more hit-and-miss when we only tested 44 molecules (**Fig. 2b**). To quantify how hit rate varies with the number experimentally tested, we randomly pulled sets of 44, 139 and 439 molecules from the 1,296, each 30 times, and asked how this affected hit rate. When only selecting 44 molecules—the number tested in the smaller library campaign—hit rates varied from 11% for one unlucky draw to 36% for a lucky one. Pulling sets of 439 molecules 30 times, the hit rate only varied from 20% to 27%. As the number of molecules experimentally tested increased, the standard deviation in hit rates decreased from 6.1% to 3.5% to 1.7% (**Fig. 2d**). This variability was even starker when plotted by affinity bin; for instance, it was not until set size rose to 439 molecules tested that the highest affinity molecules were reliably sampled (**Fig. 2e).** Re-analyzing previous campaigns against the σ2 and dopamine receptor^1,4^, where around 500 molecules were experimentally tested, similar variability was seen in both hit rates and in sampling of the high-affinity docking hits, which for σ2 were in the low nanomolar range (**Extended Data** Fig. 4).

These results suggest that both hit rates and affinities in docking screens may be unreliable when only dozens of molecules are tested, as is common in the field. To quantify how many molecules should be tested to report stable hit rates and affinity ranges, we drew on the observation that when large numbers of molecules are experimentally tested for AmpC, and for the σ2 and the dopamine D4 receptors, there is an exponential relationship between affinity and hit rate, something also seen in high-throughput screens^28^. For the top-ranking 1% of docked molecules from each campaign, we modeled hit rates (y) and hit affinities (x) with an exponential function *y* = *b*(1 – *e*^−*cx*^) for each of AmpC, σ2 and D4 (**Fig. 3a**). This functional form fit the distribution of affinities for the 1,296 molecules tested for AmpC, 327 for σ2 and 371 for D4 (all top 1% ranking molecules) with R^2^ values of 0.998, 0.998, and 0.985, respectively. As smaller sets are drawn from the full sets, variability rises (**Fig. 2**). Beginning with a sample size of 1,296, sampling was stepwise reduced by 20 molecules in a bootstrapping manner, repeating this 1,000 times to evaluate divergence (**Fig. 3b**). By ∼495 molecules, the average R^2^ of D4 curves falls to 0.95, a point on all three curves where we began to see the meaningful divergence the fit achieved over the full range of compounds plotted. This same R^2^ occurs at 215 and 135 molecules for AmpC and σ2, respectively, reflecting a relationship that is inversely proportional to the hit rate for each target among the top-scoring 1% of the docked molecules (22.4% hit rate for AmpC, 38.7% for σ2 and 20.8% for D4). In these targets, testing fewer than these several hundred compounds degrades the correlation of affinity with hit rate, which is useful for planning how many compounds should be tested. For targets with relatively high hit rates, this suggests that over a hundred molecules should be experimentally tested for confident hit rates and affinity ranges from molecular docking. For targets where one might expect lower hit rates, even more compounds would need to be tested for confident results.

**Fig. 3.**
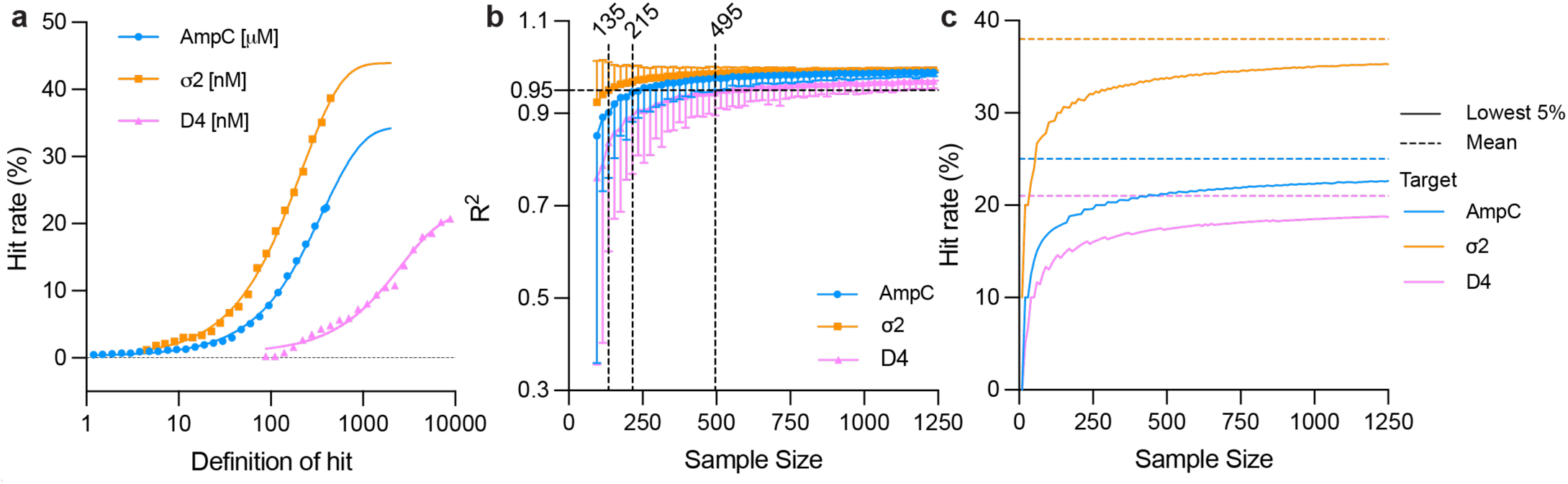
Several hundred compounds should be tested in ultra-large docking campaigns. **a,** For the top-ranking 1% of the docked molecules, the relationship between hit affinity and hit rates can be fit with an exponential plateau model *y* = *b*(1 – *e*^−*cx*^) with y represents the hit rate, x is minimum affinity to be classified as a hit (for AmpC, the unit is in micromolar and for σ2 and D4, the unit is in nanomolar), b is the maximal hit rate. The fit maximal hit rates are 34.5% for AmpC with an R^2^ of 0.998, 43% for σ2 receptor with an R^2^ of 0.998, and 20.8% for D4 with an R^2^ of 0.985. **b,** The impact of sub-sampling on the R^2^ of the fit. From among the top-ranking 1% of the docked molecules, 1,295 (AmpC, blue), 327 (σ2, orange) and 371 (D4, pink), each subsample is bootstrapped 1,000 times and fit with the parameters derived from the entire dataset. The R^2^ values are plotted with the symbols indicating the average and the error bars indicating the standard deviations of the R^2^. A dashed line of R^2^ = 0.95 is labeled. The sample sizes at which the average R^2^ value reaches 0.95 are labeled. For σ2, the sample size is 135, for AmpC, it is 215; and for D4, it is 495. **c,** Mean and 95% confidence interval for hit rate in relation to sample size for AmpC, σ2 and D4. The dashed lines show the mean hit rate from the compounds in the top 1% by docking score, and the solid line shows the boundary of a single-sided 95% confidence interval from 100,000 bootstrap iterations. Hits are defined as 400 µM affinity or better for AmpC, 677.5 nM or better for σ2 and 10 µM or better for D4.

To explore this further with a focus on hit-rate variability, we simulated random draws using the AmpC, σ2, and D4 experimental hit rates from their high-ranking compounds. One hundred thousand bootstrap iterations were performed for sample sizes ranging from 10 to 1250 compounds in increments of 10 and we considered the mean and lower bound for a single-sided 95% confidence interval at different numbers of compounds tested (**Fig. 3c**). The solid curves reflect the 95% likelihood that the hit rate will be at a certain level or higher. While the average hit rate over all simulations remains unchanged, the variability increases as the number of molecules tested drops, and so does one’s confidence that the observed hit rate reflects the true hit rate based on the overall docking rankings. This again suggests over 100 molecules may be a sensible minimum for experimental testing in large library virtual screens, even for campaigns from which one expects relatively high hit rates. Encouragingly, while both the affinity ranges and the hit rates for the screens against AmpC, σ2 and D4 differ substantially, the functional form relating hit affinity and hit number was the same and led to similar predictions for the minimum number of molecules to test for all three targets. This lends itself to predicting how many molecules would be found in different affinity ranges should one choose to test more molecules, a point to which we will return.

### Multiple novel chemotypes discovered

Only molecules topologically dissimilar from known AmpC inhibitors, and topologically diverse among each other, were selected for synthesis and testing. Since topological diversity can emerge from rearrangements and changes that leave core pharmacophores intact, we also visually inspected inhibitors for novelty. We prioritized these by two criteria: molecules that sampled new scaffolds, and molecules that explored a new warhead for binding in the crucial oxyanion recognition site of AmpC (Extended Data Fig. 5). For instance, **Z6615021877** and **Z6722203632** introduce tetrazolone and tetrazole anionic warheads, respectively, both of which were previously unknown for AmpC. **Z2607647274** and **Z4173922012** employ cycloalkyl carboxylate and tricyclo-heptane carboxylate as their warheads. Meanwhile, **Z2610488449**, which utilizes a novel urea linker scaffold, achieves a high affinity of 12 µM. The affinity of this scaffold was readily optimized to 4 µM, marking it among the most effective AmpC inhibitors that does not rely on a sulfonamide linker.

### Docking score predicts hit rate

In earlier studies against the dopamine D4 and σ2 receptors, we had found that docking score correlated to experimental hit rate, generating a well-behaved sigmoidal curve that plateaued at a maximum hit rate once more favorable docking scores were reached^1,4^. While these curves suggested an unexpected ability to categorize molecules as ligands, both receptors have unusually well-formed, buried binding sites. Moreover, the plateauing of the score vs. hit rate curve suggests a limitation in even our ability to *categorize*, far less rank-order. To investigate how docking might predict binding in a more solvent-exposed, historically more difficult binding site, we re-explored this relationship for AmpC. Docked molecules were not only selected from among the very best docking energies, as is typical in virtual screening, but also from mediocre and unfavorable docking scoring ranges. Molecules were picked from among 16 scoring “bins” beginning at the most favorable DOCK3.8 scores (-100.58 kcal/mol for AmpC) down to -28 kcal/mol. The top 1% of the docking-ranked library extends down to -72 kcal/mol scores, and ranks fall off steeply from there such that by -28 kcal/mol 49% of the 1.7 billion molecule library has been sampled. More than 50 molecules per bin were selected from the -100.58 to the -60 kcal mol^-1^ bin, and for scores worse (less negative) than -60, more than 20 molecules were tested per bin. Molecules were selected strictly by numerical rank at the beginning of each bin. These were tested for AmpC inhibition as above and docking score was plotted against experimental hit rate (number active/number tested) (**Fig. 4a**).

**Fig. 4.**
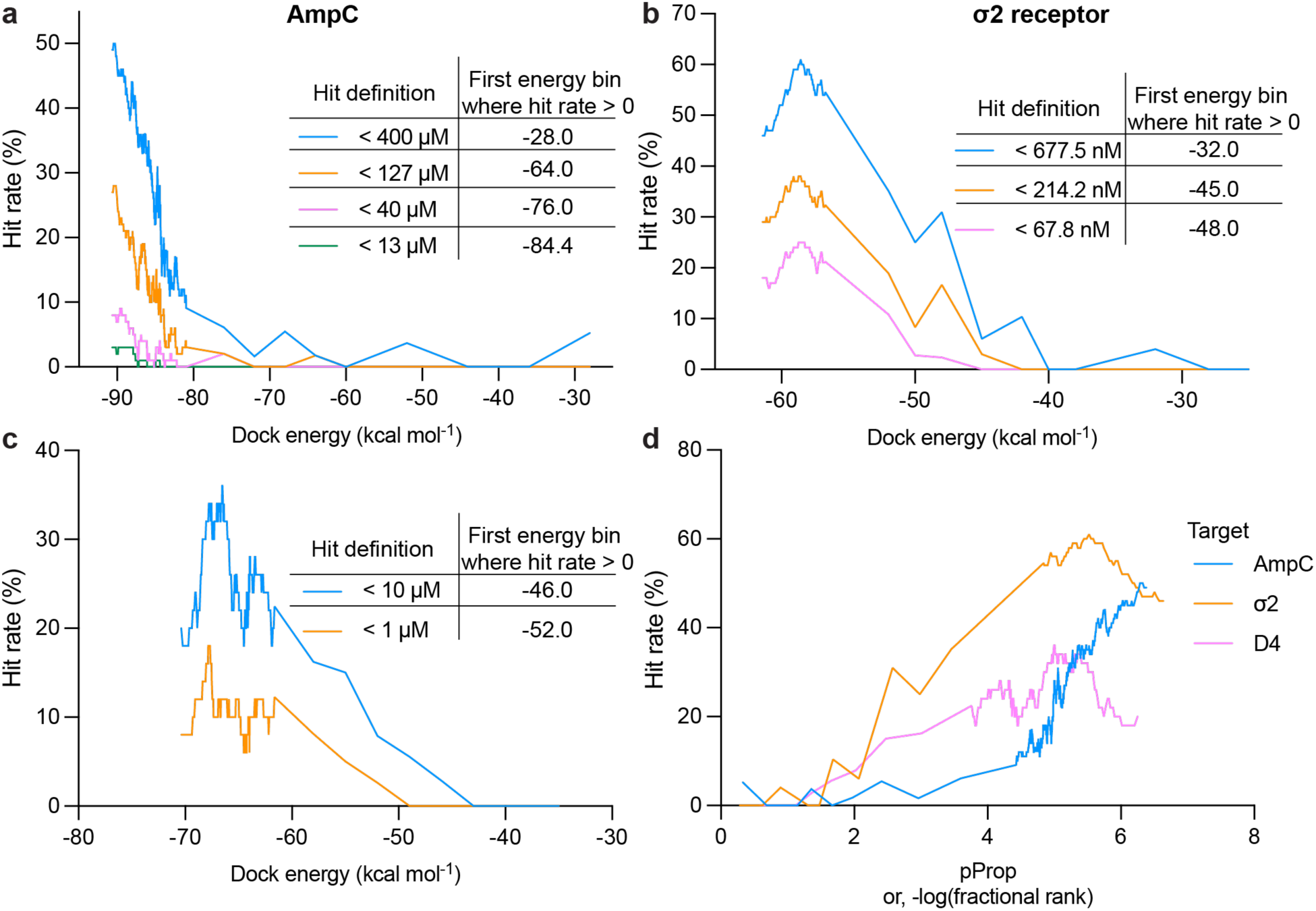
Hit rate of experimentally tested compounds plotted against DOCK scores with different affinity cutoffs. **a,** The AmpC hit rates of 1,292 auto-picked compounds using four different affinity cutoffs, < 400, 137, 40 and 13 μM, are plotted against DOCK scores. **b, σ**2 receptor hit rates of 484 compounds plotted against DOCK scores with three different affinity cutoffs: < 667.5, 241.2, 67.8 nM. **c,** Dopamine D4 hit rates of 549 compounds plotted against DOCK scores with two different affinity cutoffs: <10 and <1 μM. **d,** Rescaling the hit rate curves of the three targets by the log10 of fractional rank in the library. For each target, the most permissive hit definition is used.

Notably, hit rates fell monotonically as scores worsened (**Fig. 4a**, blue curve). This resembles what had been previously observed for the σ2 and dopamine receptors^1,4^, except that here we do ***not*** observe a hit rate plateau; hit rates begin at a maximum at the best docking scores and fall steadily as scores worsen. The difference between the AmpC results and the plateaus observed previously is that for AmpC we were careful to from the beginning exclude a small fraction of likely artifacts that concentrate among the very top scoring molecules^14^ (Extended Data Fig. 1).

To investigate how the *affinities* of docking actives also track with docking rank, we plotted docking score versus hit rate in the 400, 127, 40, and 13 µM ranges (**Fig. 4a**, orange, pink, and green curves). As for the behavior of the overall set of actives, the hit rates in each affinity-range rose steadily as score improved. Intriguingly, the more potent inhibitors appear at better scores than the less potent ones, with those in the 127 µM or better tranche beginning to appear at scores of -64, those in the 40 µM or better tranche appearing only past -76, and the most potent inhibitors only appearing at the -85 bin. This hints at docking score correlating with gross categorical ranking of affinity, something that has not apparent from smaller studies, nor even expected^29,30^. To explore generality, we undertook the same analysis with the docking campaigns against the σ2 receptor and dopamine D4 receptor, where hundreds of molecules were tested across docking ranks that ranged from high to mediocre to poor, as in this study. While the σ2 and dopamine receptor docking hits were more potent than the AmpC hits, typically in the nM range, the same patterns emerged—the most potent ligands appear at better (more negative) docking scores than did the mid-potency ligands, which appear at better scores than did those with the lowest affinity threshold (**Fig. 4b-c**). Admittedly, the relationship between docking score and affinity is mostly categorical, but it appears to rank molecules better than simple binary classification as binders or non-binders, with more potent ligands more concentrated in better-scoring regions. As loose as these correlations are, they may support a predictive relationship between docking score and affinity category (high, medium, or low), at least when tested at scale. This would warrant a renewed emphasis on improving the field’s scoring functions and offer a metric against which they might be tested.

To compare the hit rate curves for the three targets, we plotted the negative logarithm of the rank percent (“pProp”) for the dopamine and σ2 receptors, and for AmpC (**Fig. 4d**). A pProp of 3 denotes a compound occupying the top 0.1% scoring region, a pProp of 4 the top 0.01%, and so on; plotting rank avoids scoring offsets among the targets. The hit rate curve of the most permissive hit definition for each target is plotted against the pProp. The D4 and σ2 curves align well, peaking around a pProp of 5, with the plateau occupying the region from 4 to 6 (top in 10,000 to top in 1,000,000), while the AmpC curve is slightly right shifted, peaking above 6 and not suffering from a plateau. These curves allow one to quantify the parts of the docking scoring range where most hits are likely to be found. For the D4 and σ2 receptors, it also alerts one to the danger of over-emphasizing the very best ranked molecules where those that cheat the scoring function concentrate, absent controls for them^14^. As docking and virtual screening libraries climb into the tens-of-billions of molecules^5,21^, this concern will become more pressing.

## Discussion

In the last five years, the number of molecules readily accessible for ligand discovery has expanded 10,000-fold. Anecdotally, this has led to ligands with improved activity from library docking. How true this is, however, has not been quantified in apples-to-apples comparisons of smaller vs. larger libraries. Several other inferences from large library docking screens have also not been quantified, such as that testing more high-ranking molecules yields correspondingly more ligands, or that as docking score improves so too does hit rate and perhaps even affinity. Here, we begin to test these ideas experimentally; five key observations emerge^1,14^. **First,** comparing a docking screen of 99 million molecules to one of 1.7 billion molecules, against the same target, hit rates improved with library size, as did the potency of the inhibitors. Multiple new chemotypes were discovered, not previously observed as AmpC inhibitors. **Second,** consistent with the idea that there are many more ligands to be discovered than are being prioritized, the number of new inhibitors found scaled almost linearly with the number of top-ranking molecules tested; experimentally testing 30-fold more molecules led to the discovery of 50-fold more inhibitors. **Third,** to determine reliable docking statistics from a large library screen, one must also *experimentally test* at scale. When only a handful of molecules are tested, as is common in docking, statistics of hit-rates and maximal affinities will have large error ranges. This study suggests that typically several hundred molecules should be tested for docking statistics to be trustworthy. **Fourth,** in contrast to earlier studies where we saw hit rates plateau above a certain docking score^1,4^, here no plateau was observed in the hit rate vs. docking score curve—hit rates continued to climb monotonically and essentially linearly as score improves. This was also true for the dopamine and s2 receptors on re-analysis after removing their high-ranking artifacts. While more studies are necessary, this observation supports the idea that as libraries grow, hit rates and hit affinities will improve, as long as high-ranking docking artifacts can be removed or avoided. **Finally**, a loose, categorical correlation between docking score and ligand affinity was observed for AmpC, and on re-analysis also for the σ2 and dopamine D4 receptor campaigns where sufficient molecules were also tested across the scoring range to support this analysis^1,4^. While this correlation remains loose and only by relative affinity category (e.g., strong, mediocre, weak), it may suggest that further optimization of docking scoring functions will allow the field to distinguish not only binders from non-binders, but also categorically rank them by activity, something we and others have long discounted^29,30^.

Several caveats should be aired. Most importantly, the monotonic improvement of hit rate with docking score, and the loose categorical correlation between affinity and score, have only been observed in three systems. This merits investigation in other targets, ideally using other scoring functions, at scale. Current community tests of docking methods, such as CACHE^31^, may offer a forum for doing so. Methodologically, we note that for less than 10% of the molecules reported here were full IC50 curves determined. While these correlated well with inferred IC50 and Ki values based on three concentration point inhibition, such affinities must be considered approximate.

These caveats should not obscure the major observations of this study. Against the same target, docking a 20-fold larger library led to improved hit rates and affinities, consistent with theoretical simulations^14^. Similarly, as more high-ranking molecules are tested, more ligands are found, supporting the idea that most true ligands in the new ultra-large libraries remain to be tested (we suffer from an embarrassment of riches). Once we correct for high-ranking docking artifacts, hit rate rise monotonically with docking score. More tentatively, a correlation between affinity and score also appears at scale. How this holds up will depend on further testing, but even now these results support continued investment in library growth and methods that can exploit it. While brute force docking, of the sort described here, has been able to address a 1000-fold increase in library size, to go up another thousand-fold, into the trillions of molecules, seems beyond it, and more guided sampling of chemical space may be required^5,11,32,33^ To support such efforts, we are making available the identity, docking scores, and experimental activities of each of the 1521 molecules tested in the enzyme assay (**SI Table 1**), and extensive docking score and pose information from the full library screen (https://lsd.docking.org). What this study does suggest is that efforts to sample from the supra-trillion molecule space should be worthwhile.

## Data and code availability

The compounds docked in this study are freely available from the ZINC20 and ZINC22 databases, https://zinc20.docking.org and https://cartblanche22.docking.org. All compounds tested can be purchased from Enamine. Compound information including their ZINC ID, SMILES, DOCK score, ranking, and affinity can be found in **SI Table 1**. The synthetic procedures and purity information for the hits can be found in the **SI Data 2** and **SI Table 3**. Extensive docking-related files can be found at https://lsd.docking.org. DOCK3.8 is freely available for non-commercial research at https://dock.compbio.ucsf.edu/DOCK3.8/. A web-based version is available without restriction at https://blaster.docking.org/. X-ray structures and maps are available in the Protein Data Bank under accession numbers PDBID 9C81 (Z4462773688), PDBID 9C6P (Z6615017509), PDBID 9C83 (Z8427841182), PDBID 9C84 (Z6615020275), and PDBID 9C8J (Z6615017782) respectively.

## Acknowledgements

Supported by US National Institutes of Health (NIH) grants R35GM122481 (to BKS) and GM71896 (to JJI), and a Damon Runyon Postdoctoral Research Fellowship (FL). We thank ChemAxon for JChem, OpenEye Scientific software for OEChem and Omega2, Molecular Networks for Corina, Molinspriation for Mitools, and Schrodinger LLC for the Maestro suite. We thank George Meigs and James Holton for their assistance at Beamline 8.3.1 at the Advanced Light Source, operated by UCSF with NIH grants R01 GM124149 for technology development and P30 GM124169 for beamline operations, and the Integrated Diffraction Analysis Technologies program of the US Department of Energy Office of Biological and Environmental Research. We thank Karthik Srinivasan for his assistance on data collection. The Advanced Light Source (Berkeley, CA) is a national user facility operated by Lawrence Berkeley National Laboratory on behalf of the US Department of Energy under contract number DE-AC02-05CH11231, Office of Basic Energy Sciences.

## Author Contributions

FL conducted the docking screens and the ligand optimization assisted by SV and advised by BKS. FL and IG conducted the *in vitro* enzymatic assays, with early assistance from SV. FL determined the structures by X-ray crystallography, with assistance from VB, XX, advised by JF. FL and OM did the analysis with advice from MS. Aggregation studies were conducted by KFV and IG. JJI developed and prepared the make-on-demand library assisted with large library docking strategies. DSR and YSM supervised compound synthesis of Enamine compounds purchased from the ZINC22 database and the 46 billion catalog library.

## Methods

### Large-scale docking

The campaign used the structure in PDB 1L2S. Three Q120 conformations were modeled based on the X-ray density of 3FKW using qFit-3.0, with an occupancy of 0.49, 0.34 and 0.17^34^. The occupancy of the alternative conformations was converted into an additional energy term and incorporated in the DOCK scoring function as described previously^35^. The protein structure was protonated using Reduce^36^. Energy grids for the different energy terms of the scoring function were pre-generated--van der Waals term based on the AMBER force fields using CHEMGRID^37^; Poisson–Boltzmann-based electrostatic potentials using QNIFFT73^38,39^; context-dependent ligand desolvation was calculated using SOLVMAP^40^. The volume of the low dielectric and the desolvation volume was extended out 2.0 and 0.25 Å. The thiophene carboxylate inhibitor solved in PDB 1L2S was used to generate matching spheres, which are later used by the docking software to fit pre-generated ligands’ conformations into the small molecule binding sites^41^. The resulting docking set-ups were evaluated for its ability to enrich known AmpC ligands over property-matched decoys. Decoys are theoretical non-binders to the receptor as they are topologically dissimilar to known ligands but retain similar physical properties. We curated 31 AmpC ligands based on their dissimilarity among themselves. 2,480 decoys were generated by using the DUDE-Z pipeline^42^. The docking set-up can rank ligands over decoys with a logAUC of 28.5 with the majority of the ligands recapitulating their experimental poses. For docking against 1.7-billion molecules, each molecule from the ZINC22 database^43^ was sampled in about 3,822 orientations and 875 conformations by using DOCK3.8^41^. Overall, over 1841 trillion complexes were sampled and scored, spending 2,129,230 core hours, or about a month on a 3,000 core cluster, using DOCK3.8^41^.

### Hit-picking strategy

To increase novelty, high-ranking molecules with scores down to -79.25 (99,277 molecules), and molecules from different energy bins (25,000 from -76, -72, -68, -64 and -60 bins and 5,000 from -52, -44, -36 and -28 bins), summed to 244,277 molecules, were filtered to exclude those similar to 237 previously known ligands. A Tc cutoff of 0.5 was used; no molecule more similar than this value was allowed, removing 9,712 molecules. We also filtered out molecules that buried too many uncompensated polar groups—while DOCK3.8 penalizes for desolvation, we find that these artifacts can nevertheless occur. Using LUNA 1,024-length binary fingerprints^23^, molecules that had more than 1 hydrogen bond donor and more than 6 hydrogen bond acceptors that were not compensated with polar interactions to the protein were removed; 40,687 molecules were filtered-out at this step. This left 193,878 for further processing. For autopicking, these molecules were clustered for self-similarity using an ECFP4 *Tc* = 0.32, resulting in 80,767 cluster heads.

Most of the molecules tested were “autopicked” based on docking rank. With almost all of the high-ranking molecules being negatively charged, we wanted to ensure that their representation as anions at pH 7.4 was likely. We used JChem to calculate the distribution of protonation states of the high-ranking cluster heads and compared this to the dominant state represented in their docked poses (multiple protonation states of a molecule can be docked). Only when the calculated dominant charge state matched with that of the docked pose, and the species is calculated to be more <u>than</u> 80% anionic, was the molecule accepted for autopicking, which left 56,814 molecules. Molecules were picked based on their docking ranks across different affinity bins, selecting 1,274 molecules for synthesis and testing.

For manual picking from the different energy bins, all cluster heads were again filtered for interactions using LUNA, seeking molecules that formed hydrogen bonds with backbone of A318, that made pi-pi interactions with Y221, and that made at least two more interactions with the binding pocket (i.e. hydrogen bonds with N152, N346, G320, S212, R204, Q120, cation-pi with K315, K67, or pi-pi interaction with Y150). The molecules that passed these filters were reclustered at a *Tc* = 0.32; cluster heads were visually inspected and prioritized. The rest of the high-scoring cluster heads were also manually inspected seeking new interesting chemotypes. A total of 687 were prioritized manually, slightly less than half of the molecules that were synthesized and tested.

### AmpC enzymology

All candidate inhibitors were dissolved in DMSO at 20 mM, and more dilute DMSO stocks were prepared as necessary so that the concentration of DMSO was held constant at 1% v/v in 50 mM sodium cacodylate buffer, pH 6.5. AmpC activity and inhibition was monitored spectrophotometrically using either CENTA or nitrocefin as substrates. All assays included 0.01% Triton X-100 to reduce compound aggregation artifacts. Active compounds were further investigated for aggregation by dynamic light scattering (DLS) and by detergent-dependent inhibition of the counter-screening enzyme malate dehydrogenase.

For initial screening, the docking hits were diluted such that final concentrations in the reaction buffer was 200 μM, 100 μM, and 40 μM. In these assays, two widely-studied AmpC substrates were used, depending on availability, CENTA^44^ and nitrocefin^16^. The first was tested at an [S]/Km ration of 1.81 (Km CENTA 27.6 µM; [S] = 50 µM) and the second was tested [S]/Km ratios of 0.556 (Km nitrocefin 180 µM; [S] = 100 µM) and 0.156 ([S] = 28 µM). The colorimetric assay was converted to a medium throughput manner using a BMG Labtech CLARIOstar. Substrate (CENTA (EC50 = 27.6 μM) or nitrocefin (EC50 = 180 μM)) and protein were injected into buffer containing the putative inhibitor, followed by rate measurement for 50 seconds in 96-well format. IC50 values reflect the percentage inhibition fit to a dose-response equation in GraphPad Prism with a Hill coefficient set to one 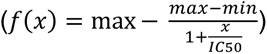. The Ki was calculated using the Cheng-Prusoff equation 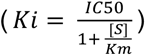. For 18 of the more potent compounds, based on the initial three concentration-point results, full dose response curves were measured, and for another eight full Ki values were measured and calculated using Lineweaver-Burk plots.

### AmpC crystallization, data collection and structure determination

AmpC crystallization was carried out as previously described^16^. Briefly, co-crystals of AmpC and inhibitors were grown by vapor diffusion in hanging drops equilibrated over 1.7 M potassium phosphate buffer (pH 8.7) using microseeding. The initial concentration of protein in the drop was 6 mg/mL and the concentration of the inhibitor was 0.5 mM. The inhibitor was added to the crystallization drop in a 4% DMSO, 1.7 M potassium phosphate buffer (pH 8.7) solution. Crystals appeared within 3–5 days after equilibration at 23°C.

Data were measured from a single crystal per complex on the Beamline 8.3.1 of the Berkeley Advanced Light Source, with wavelength 1.11583 Å at 100 K. Before data collection, co-crystals of AmpC were immersed in a cryoprotectant solution of 20% sucrose, and 1.7 M potassium phosphate (pH 8.7) for about 20 s and then flash-cooled in liquid nitrogen. The structures were solved by molecular replacement with PHENIX^45^ using PDB 1L2S as the search model. Structure refinement was carried out with PHENIX and COOT^46^. MolProbity^47^ was used for validation (**Extended Data Table 3**); structural figures were prepared using ChimeraX^48^.

### Hit rate curves

To obtain hit rate curves, the experimentally tested molecules for each target (AmpC, the σ2 and dopamine D4 receptors) were ordered by increasing DOCK score. A rolling window was passed over the list, calculating the hit rate as the percentage of molecules with experimentally determined affinity equal to or better than the hit definition, and the DOCK score as the average for the window. A window size of 100 was used for AmpC and σ2, and a window of 50 for D4 receptor. For all three targets, molecules were picked from both within and outside of what would typically be considered high-ranking regions. The rolling window was stopped for those scores outside the high-ranking region since discrete score bins were used in the hit-picking of these likely non-binders. The scores at which the rolling window was stopped are -78 for AmpC, -52.5 for σ2 and -60 for D4. For the pProp rescaling, the same strategy was used, but the DOCK scores were transformed to fractional rank based on the observed score distribution. The negative base 10 logarithm of the fractional rank is then reported, termed “pProp”.

### Hit rate modelling

For sampling hit rate variability in relation to sample size, we used sample sizes for 10 to 1250 in jumps of 10. For each sample size, we picked 100,000 random samples of the uniform distribution [0, 1]. The hit rate of the sample was then defined as the number of observations with equal to or lower than the observed experimental hit rate for that target. A single-sided 95% confidence interval is built by taking the boundary value between the top 95% observed hit rates and the bottom 5%.

## References

1 Lyu, J. et al. Ultra-large library docking for discovering new chemotypes. Nature 566, 224–229, doi:10.1038/s41586-019-0917-9 (2019).

2 Gorgulla, C. et al. An open-source drug discovery platform enables ultra-large virtual screens. Nature 580, 663–668, doi:10.1038/s41586-020-2117-z (2020).

3 Stein, R. M. et al. Virtual discovery of melatonin receptor ligands to modulate circadian rhythms. Nature 579, 609–614, doi:10.1038/s41586-020-2027-0 (2020).

4 Alon, A. et al. Structures of the sigma(2) receptor enable docking for bioactive ligand discovery. Nature 600, 759–764, doi:10.1038/s41586-021-04175-x (2021).

5 Sadybekov, A. A. et al. Synthon-based ligand discovery in virtual libraries of over 11 billion compounds. Nature 601, 452–459, doi:10.1038/s41586-021-04220-9 (2022).

6 Fink, E. A. et al. Structure-based discovery of nonopioid analgesics acting through the alpha(2A)-adrenergic receptor. Science 377, eabn7065, doi:10.1126/science.abn7065 (2022).

7 Singh, I. et al. Structure-based discovery of conformationally selective inhibitors of the serotonin transporter. Cell 186, 2160–2175 e2117, doi:10.1016/j.cell.2023.04.010 (2023).

8 Gahbauer, S. et al. Docking for EP4R antagonists active against inflammatory pain. Nat Commun 14, 8067, doi:10.1038/s41467-023-43506-6 (2023).

9 Sadybekov, A. V. & Katritch, V. Computational approaches streamlining drug discovery. Nature 616, 673–685, doi:10.1038/s41586-023-05905-z (2023).

10 Gorgulla, C. et al. A multi-pronged approach targeting SARS-CoV-2 proteins using ultra-large virtual screening. iScience 24, 102021, doi:10.1016/j.isci.2020.102021 (2021).

11 Klarich, K., Goldman, B., Kramer, T., Riley, P. & Walters, W. P. Thompson Sampling horizontal line An Efficient Method for Searching Ultralarge Synthesis on Demand Databases. J Chem Inf Model 64, 1158–1171, doi:10.1021/acs.jcim.3c01790 (2024).

12 Walters, W. P. Virtual Chemical Libraries. J Med Chem 62, 1116–1124, doi:10.1021/acs.jmedchem.8b01048 (2019).

13 Gorgulla, C., Jayaraj, A., Fackeldey, K. & Arthanari, H. Emerging frontiers in virtual drug discovery: From quantum mechanical methods to deep learning approaches. Curr Opin Chem Biol 69, 102156, doi:10.1016/j.cbpa.2022.102156 (2022).

14 Lyu, J., Irwin, J. J. & Shoichet, B. K. Modeling the expansion of virtual screening libraries. Nat Chem Biol 19, 712–718, doi:10.1038/s41589-022-01234-w (2023).

15 Weston, G. S., Blazquez, J., Baquero, F. & Shoichet, B. K. Structure-based enhancement of boronic acid-based inhibitors of AmpC beta-lactamase. J Med Chem 41, 4577–4586, doi:10.1021/jm980343w (1998).

16 Powers, R. A., Morandi, F. & Shoichet, B. K. Structure-based discovery of a novel, noncovalent inhibitor of AmpC beta-lactamase. Structure 10, 1013–1023, doi:10.1016/s0969-2126(02)00799-2 (2002).

17 Feng, B. Y., Shelat, A., Doman, T. N., Guy, R. K. & Shoichet, B. K. High-throughput assays for promiscuous inhibitors. Nat Chem Biol 1, 146–148, doi:10.1038/nchembio718 (2005).

18 Feng, B. Y. et al. A high-throughput screen for aggregation-based inhibition in a large compound library. J Med Chem 50, 2385–2390, doi:10.1021/jm061317y (2007).

19 Eidam, O. et al. Design, synthesis, crystal structures, and antimicrobial activity of sulfonamide boronic acids as beta-lactamase inhibitors. J Med Chem 53, 7852–7863, doi:10.1021/jm101015z (2010).

20 Babaoglu, K. et al. Comprehensive mechanistic analysis of hits from high-throughput and docking screens against beta-lactamase. J Med Chem 51, 2502–2511, doi:10.1021/jm701500e (2008).

21 Gorgulla, C. et al. VirtualFlow 2.0 - The Next Generation Drug Discovery Platform Enabling Adaptive Screens of 69 Billion Molecules. bioRxiv, 2023.2004.2025.537981, doi:10.1101/2023.04.25.537981 (2023).

22 Bender, B. J. et al. A practical guide to large-scale docking. Nat Protoc 16, 4799–4832, doi:10.1038/s41596-021-00597-z (2021).

23 Fassio, A. V. et al. Prioritizing Virtual Screening with Interpretable Interaction Fingerprints. J Chem Inf Model 62, 4300–4318, doi:10.1021/acs.jcim.2c00695 (2022).

24 Cheng, Y. & Prusoff, W. H. Relationship between the inhibition constant (K1) and the concentration of inhibitor which causes 50 per cent inhibition (I50) of an enzymatic reaction. Biochem Pharmacol 22, 3099–3108, doi:10.1016/0006-2952(73)90196-2 (1973).

25 McGovern, S. L., Helfand, B. T., Feng, B. & Shoichet, B. K. A specific mechanism of nonspecific inhibition. J Med Chem 46, 4265–4272, doi:10.1021/jm030266r (2003).

26 Feng, B. Y. & Shoichet, B. K. A detergent-based assay for the detection of promiscuous inhibitors. Nat Protoc 1, 550–553, doi:10.1038/nprot.2006.77 (2006).

27 O’Donnell, H. R., Tummino, T. A., Bardine, C., Craik, C. S. & Shoichet, B. K. Colloidal Aggregators in Biochemical SARS-CoV-2 Repurposing Screens. J Med Chem 64, 17530–17539, doi:10.1021/acs.jmedchem.1c01547 (2021).

28 Walters, W. P. & Namchuk, M. Designing screens: how to make your hits a hit. Nat Rev Drug Discov 2, 259–266, doi:10.1038/nrd1063 (2003).

29 Tirado-Rives, J. & Jorgensen, W. L. Contribution of conformer focusing to the uncertainty in predicting free energies for protein-ligand binding. J Med Chem 49, 5880–5884, doi:10.1021/jm060763i (2006).

30 Irwin, J. J. & Shoichet, B. K. Docking Screens for Novel Ligands Conferring New Biology. J Med Chem 59, 4103–4120, doi:10.1021/acs.jmedchem.5b02008 (2016).

31 Ackloo, S. et al. CACHE (Critical Assessment of Computational Hit-finding Experiments): A public-private partnership benchmarking initiative to enable the development of computational methods for hit-finding. Nat Rev Chem 6, 287–295, doi:10.1038/s41570-022-00363-z (2022).

32 Gentile, F. et al. Artificial intelligence-enabled virtual screening of ultra-large chemical libraries with deep docking. Nat Protoc 17, 672–697, doi:10.1038/s41596-021-00659-2 (2022).

33 Yang, Y. et al. Efficient Exploration of Chemical Space with Docking and Deep Learning. J Chem Theory Comput 17, 7106–7119, doi:10.1021/acs.jctc.1c00810 (2021).

34 Riley, B. T. et al. qFit 3: Protein and ligand multiconformer modeling for X-ray crystallographic and single-particle cryo-EM density maps. Protein Sci 30, 270–285, doi:10.1002/pro.4001 (2021).

35 Fischer, M., Coleman, R. G., Fraser, J. S. & Shoichet, B. K. Incorporation of protein flexibility and conformational energy penalties in docking screens to improve ligand discovery. Nat Chem 6, 575–583, doi:10.1038/nchem.1954 (2014).

36 Word, J. M., Lovell, S. C., Richardson, J. S. & Richardson, D. C. Asparagine and glutamine: using hydrogen atom contacts in the choice of side-chain amide orientation. J Mol Biol 285, 1735–1747, doi:10.1006/jmbi.1998.2401 (1999).

37 Meng, E. C., Shoichet, B. K. & Kuntz, I. D. Automated Docking with Grid-Based Energy Evaluation. J Comput Chem 13, 505–524, doi:DOI 10.1002/jcc.540130412 (1992).

38 Gallagher, K. & Sharp, K. Electrostatic contributions to heat capacity changes of DNA-ligand binding. Biophys J 75, 769–776, doi:10.1016/S0006-3495(98)77566-6 (1998).

39 Sharp, K. A. Polyelectrolyte Electrostatics - Salt Dependence, Entropic, and Enthalpic Contributions to Free-Energy in the Nonlinear Poisson-Boltzmann Model. Biopolymers 36, 227–243, doi:DOI 10.1002/bip.360360210 (1995).

40 Mysinger, M. M. & Shoichet, B. K. Rapid context-dependent ligand desolvation in molecular docking. J Chem Inf Model 50, 1561–1573, doi:10.1021/ci100214a (2010).

41 Coleman, R. G., Carchia, M., Sterling, T., Irwin, J. J. & Shoichet, B. K. Ligand pose and orientational sampling in molecular docking. PLoS One 8, e75992, doi:10.1371/journal.pone.0075992 (2013).

42 Stein, R. M. et al. Property-Unmatched Decoys in Docking Benchmarks. J Chem Inf Model 61, 699–714, doi:10.1021/acs.jcim.0c00598 (2021).

43 Tingle, B. I. et al. ZINC-22 horizontal line A Free Multi-Billion-Scale Database of Tangible Compounds for Ligand Discovery. J Chem Inf Model 63, 1166–1176, doi:10.1021/acs.jcim.2c01253 (2023).

44 Eidam, O. et al. Fragment-guided design of subnanomolar beta-lactamase inhibitors active in vivo. Proc Natl Acad Sci U S A 109, 17448–17453, doi:10.1073/pnas.1208337109 (2012).

45 Liebschner, D. et al. Macromolecular structure determination using X-rays, neutrons and electrons: recent developments in Phenix. Acta Crystallogr D Struct Biol 75, 861–877, doi:10.1107/S2059798319011471 (2019).

46 Emsley, P., Lohkamp, B., Scott, W. G. & Cowtan, K. Features and development of Coot. Acta Crystallogr D Biol Crystallogr 66, 486–501, doi:10.1107/S0907444910007493 (2010).

47 Chen, V. B. et al. MolProbity: all-atom structure validation for macromolecular crystallography. Acta Crystallogr D Biol Crystallogr 66, 12–21, doi:10.1107/S0907444909042073 (2010).

48 Pettersen, E. F. et al. UCSF ChimeraX: Structure visualization for researchers, educators, and developers. Protein Sci 30, 70–82, doi:10.1002/pro.3943 (2021).

