## Supplementary materials for "The impact of Library Size and Scale of Testing on Virtual Screening"

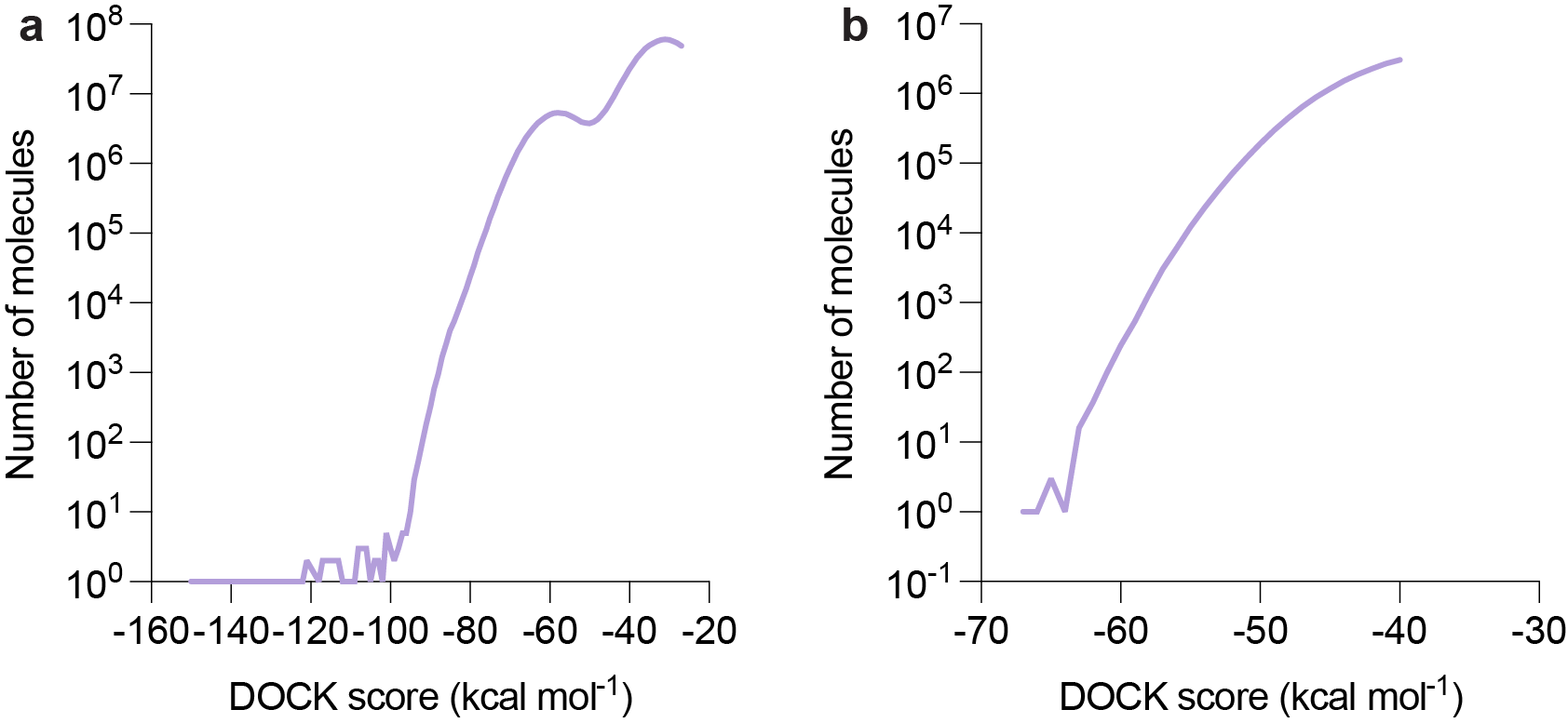

**Extended Data Fig. 1: Molecules with artifactually favorable scores disrupt the distribution of docking scores and concentrate among the top-ranking docked molecules for AmpC (a) and for the σ2 receptor^1^ (b).**

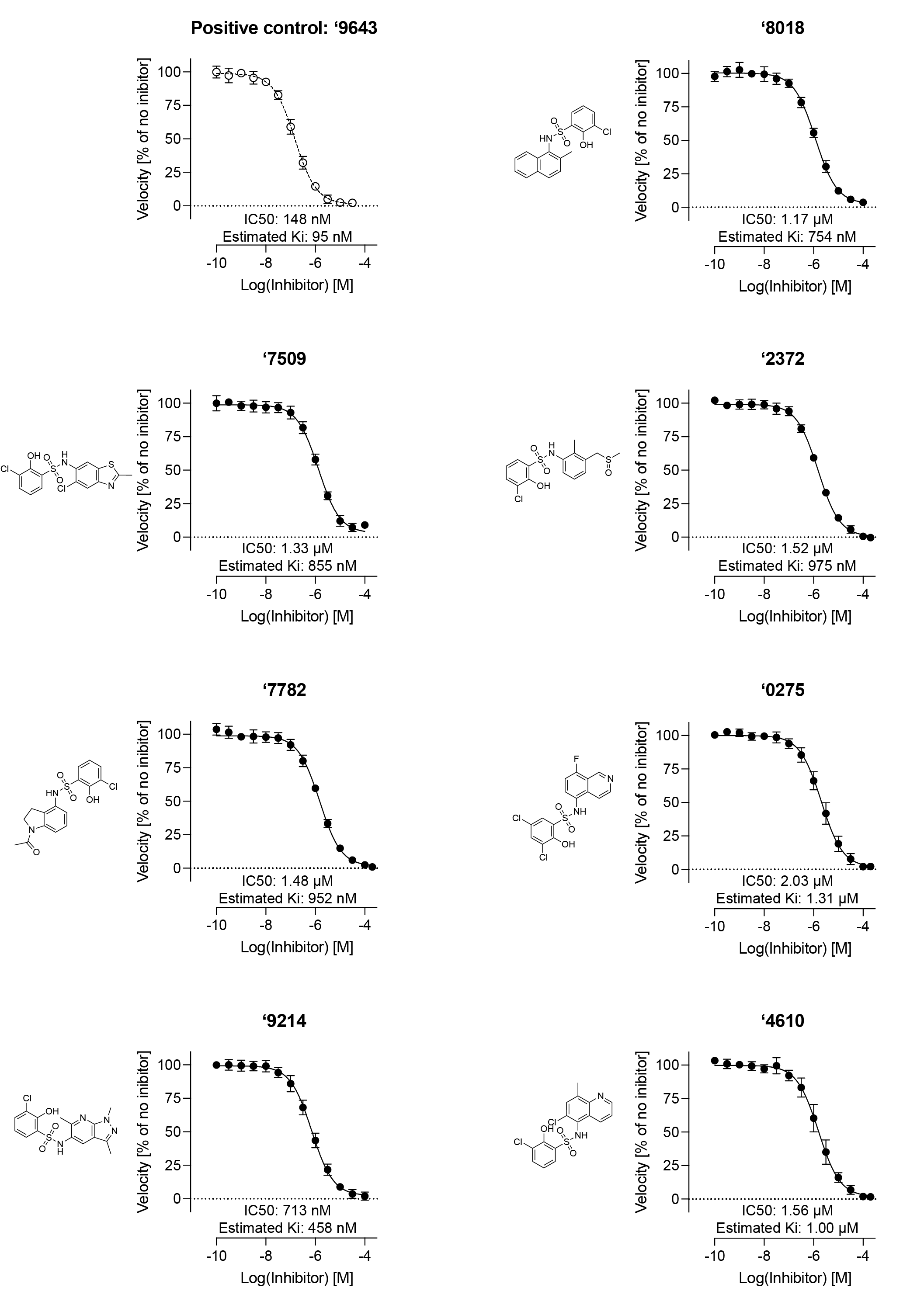

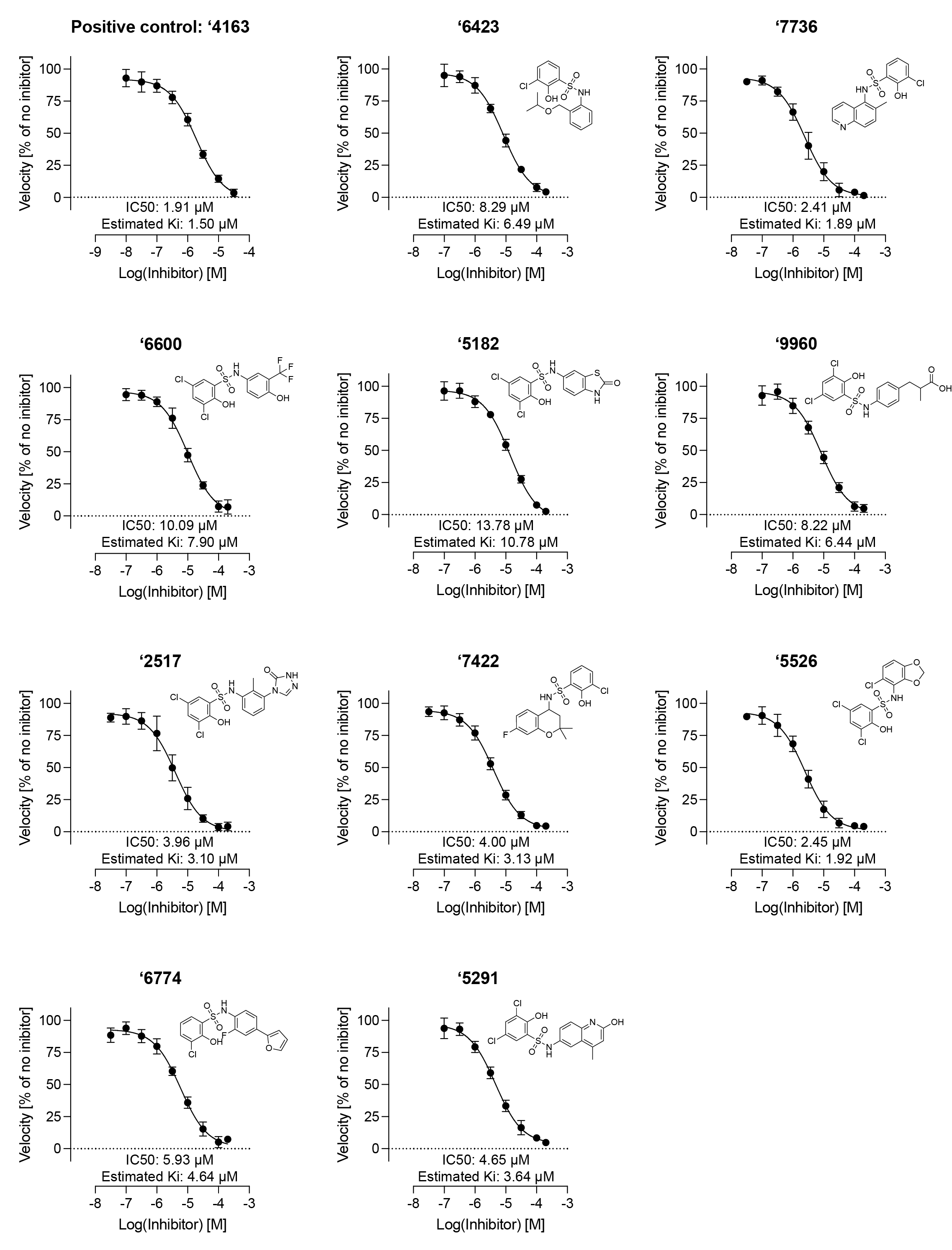

**Extended Data Fig. 2: Concentration-response curves for 17 of the new docking-derived AmpC inhibitors.** Nitrocefin was kept at a constant concentration of 100 μM (for positive control **‘9643**, new inhibitors **‘8018**, **‘7509**, **‘2372**, **‘7782**, **‘0275**, **‘9214** and **‘4610**) or 50 μM (for positive control **‘4163**, new inhibitors **‘6423**, **‘7736**, **‘6600**, **‘5182**, **‘9960**, **‘2517**, **‘7422**, **‘5526**, **‘6774** and **‘5291**). The estimated Ki is calculated based on the K_d_ of nitrocefin (180 μM) calculated from a Lineweaver-Burk analysis. The previously reported K_i_ for **‘9643** is 77 nM^2^ and for **‘4163** is 1.25 μM^2^. Data represent mean and standard deviations from three biological replicates.

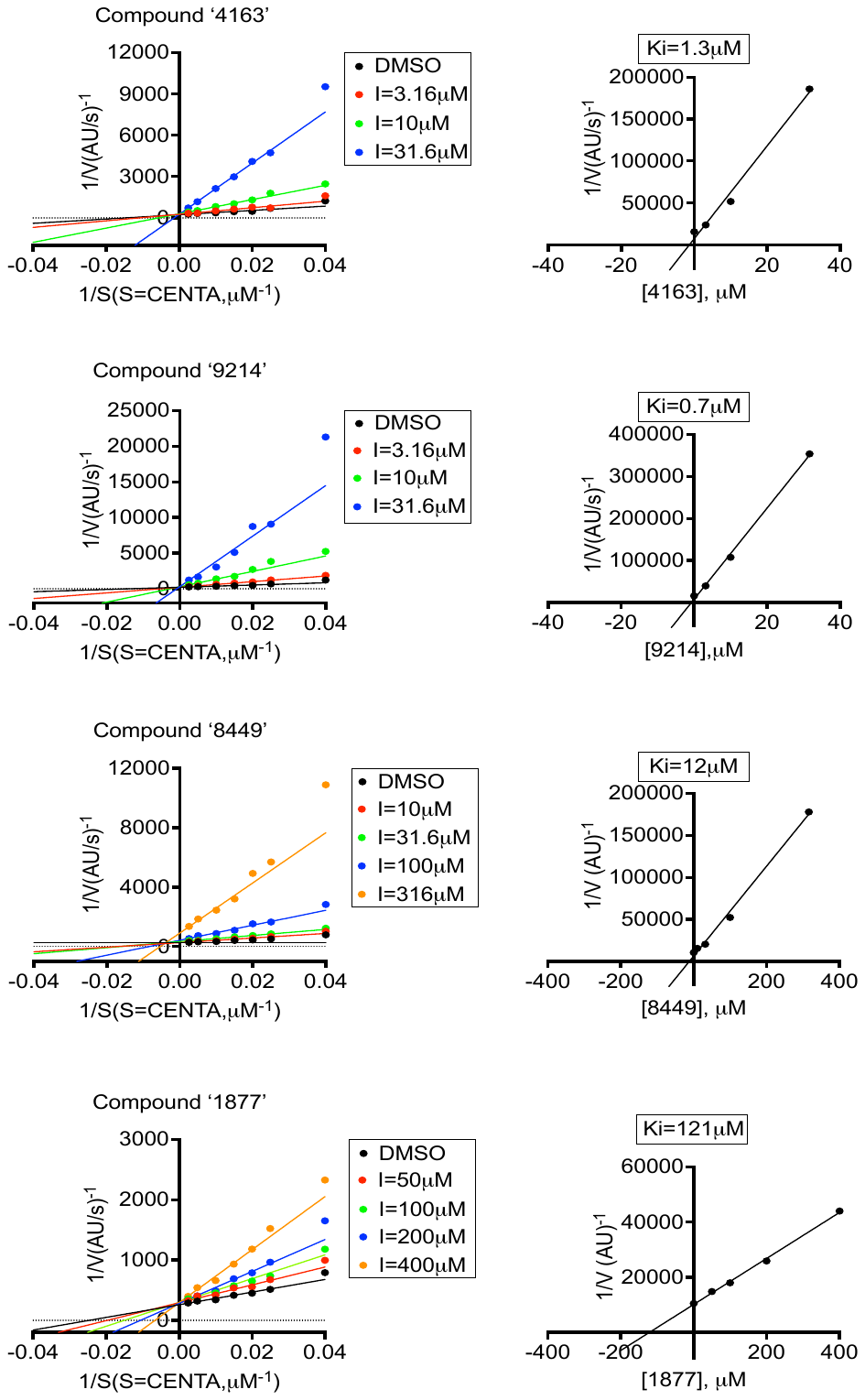

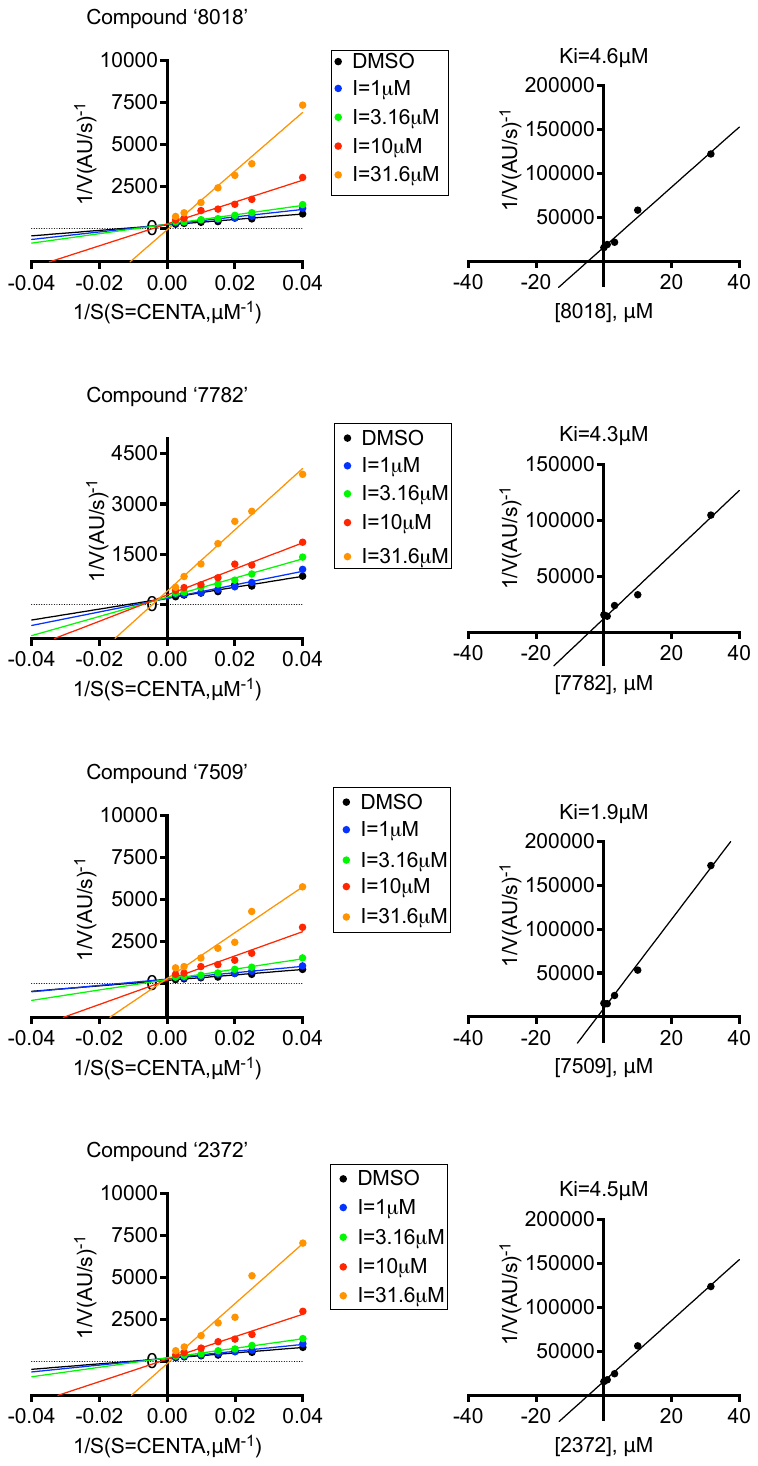

**Extended Data Fig. 3: Lineweaver-Burk plots of seven of the new AmpC inhibitors. ‘4163** is a positive control inhibitor identified in a previous docking campaign^2^.

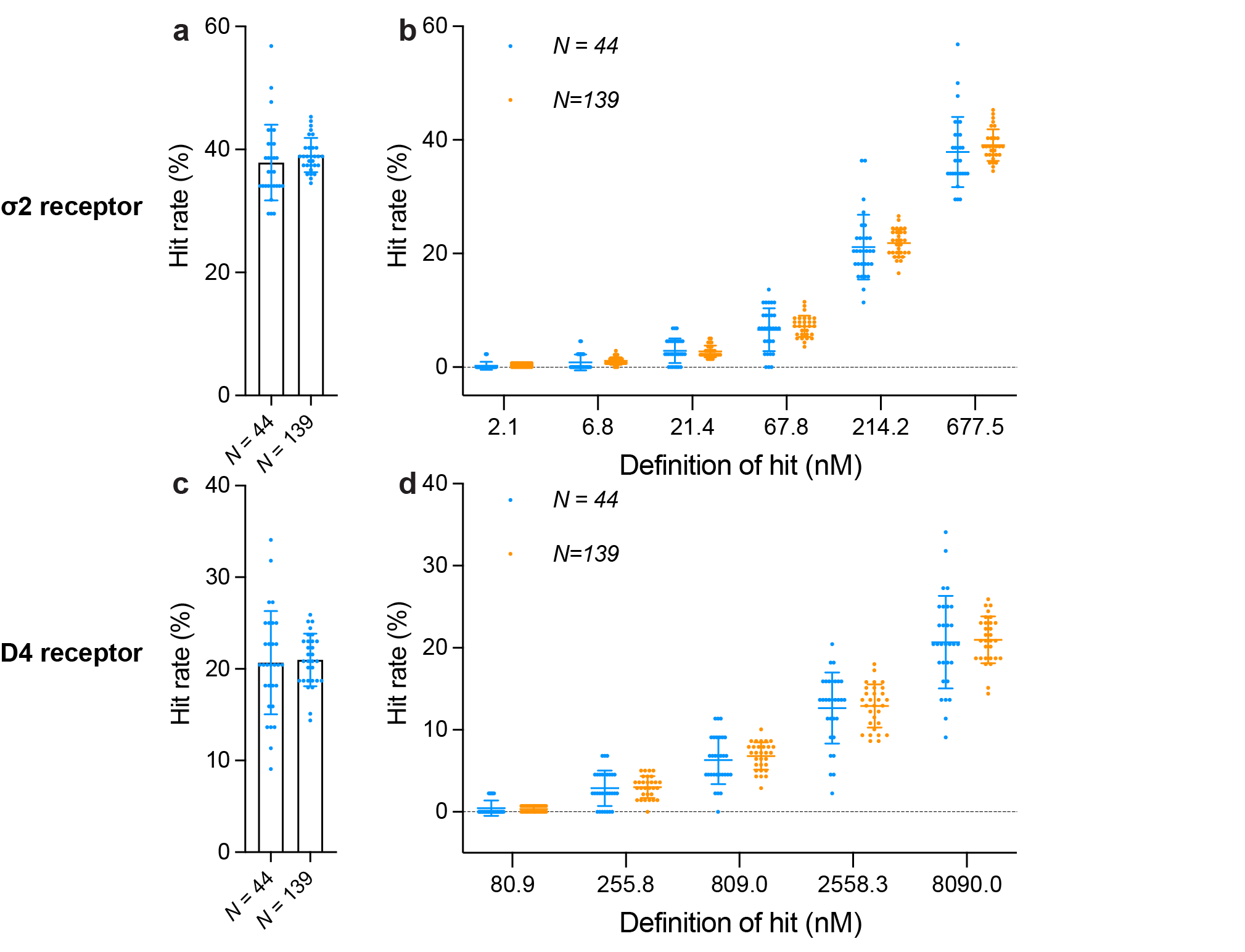

**Extended Data Fig. 4:** **The impact of testing fewer molecules on hit rate confidence.** **a**. For 327 molecules tested against the σ2 receptor, each sample size is randomly drawn 30 times and the resulting hit rates were plotted. The error bars represent SDs of the hit rates. **b**. The impact of randomly purchasing 44 and 139 molecules out of 327 molecules for testing on hit rates with different affinity cutoffs. Each sample size is drawn 30 times and the resulting hit rates were plotted. The error bars represent SDs of the hit rates. **c**. For **371** molecules tested against the D4 receptor, each sample size is randomly drawn 30 times and the resulting hit rates were plotted. The error bars represent SDs of the hit rates. **d**. The impact of randomly purchasing 44 and 139 molecules out of 371 molecules for testing on hit rates with different affinity cutoffs. Each sample size is drawn 30 times and the resulting hit rates were plotted. The error bars represent SDs of the hit rates.

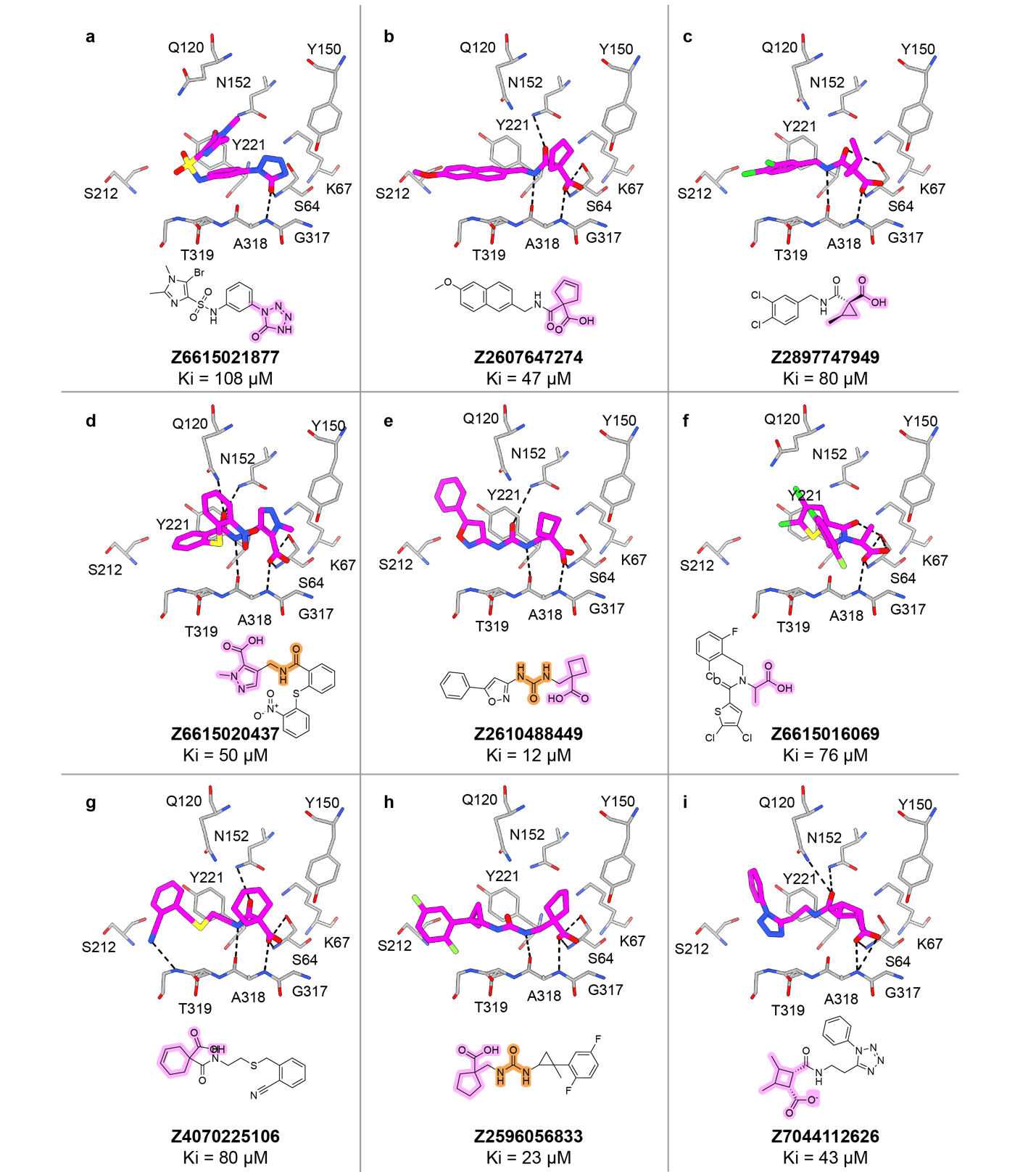

**Extended Data Fig. 5: Examples of the new warheads and chemotypes from the AmpC screen, in their docked poses in the enzyme active site.**

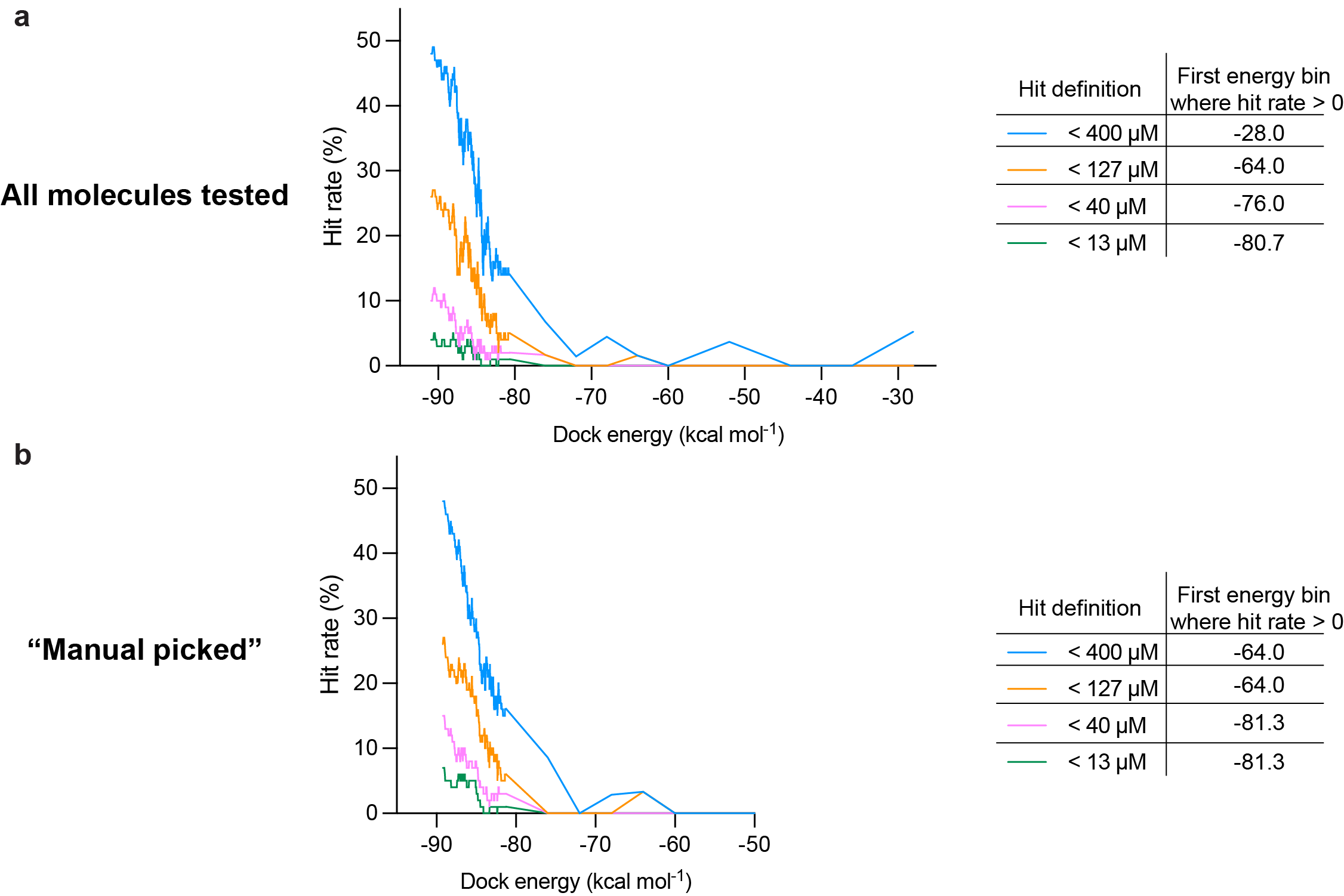

**Extended Data Fig. 6:** **Hit rate of experimentally tested compounds plotted against DOCK scores with different affinity cutoffs.** **a,** Hit rate of all compounds tested (1,447 well-behaved molecules among 1,521 purchased) plotted against DOCK scores with four different affinity cutoffs: < 400, <137, <40 and <13 μM. **b,** Hit rate of manually picked compounds (687 compounds) plotted against DOCK scores with four different affinity cutoffs: <400, <137, <40 and <13 μM.

**Extended Data Table 1. Comparison of Ki derived from 3-point inhibition and full dose response curves.**

| ZINC ID | Ki derived from 3-point inhibition | Ki derived from dose response curves |
| --- | --- | --- |
| Z2275216423 | 11.47 | 6.49 |
| Z6615017736 | 3.49 | 1.89 |
| Z2940316600 | 8.13 | 7.90 |
| Z2940315182 | 10.75 | 10.78 |
| Z6615019960 | 11.04 | 6.44 |
| Z2940322517 | 7.45 | 3.10 |
| Z6615017422 | 4.91 | 3.13 |
| Z6615015266 | 5.60 | 1.92 |
| Z6615016774 | 7.08 | 4.64 |
| Z6615155291 | 7.33 | 3.64 |

**Extended Data Table 2. Aggregation assays of selected AmpC inhibitors.**

| ZINC ID | Vendor ID | Mean DLS intensity (Cnts/s) | SD DLS intensity (Cnts/s) | Mean MDH activity (%) | SD AmpC activity |
| --- | --- | --- | --- | --- | --- |
| ZINCoI0000041Pj9 | Z6615018376 | 15,550,254 | 2,470,781 | 78 | 13 |
| ZINCoB000004161h | Z6615019227 | 1,141,892 | 161,503 | 79 | 10 |
| ZINCpH000009MPuP* | Z6615017422 | 4,840,558 | 818,412 | 102 | 20 |
| ZINCop0000098EyS | Z6615014745 | 435,473 | 80,963 | 103 | 27 |
| ZINCpO00000dcctN | Z2940308988 | 34,798,844 | 4,926,367 | 110 | 21 |
| ZINCnf0000018phh | Z2721472505 | 561,071 | 264,820 | 133 | 6 |
| ZINCnx000002khQw | Z6615020415 | 458,466 | 78,123 | 78 | 21 |
| ZINCt500000ru1Ns | Z6615016818 | 759,481 | 388,465 | 90 | 15 |
| ZINCph00000FiMLj | Z6615018469 | 104,409,329 | 5,003,599 | 137 | 11 |
| ZINCnB000000QhkZ | Z6615016830 | 572,823 | 122,079 | 76 | 23 |
| ZINCpw0000078zSl | Z2940311870 | 28,622,119 | 3,168,302 | 98 | 25 |
| ZINCon00000986RR | Z6615015390 | 543,346 | 116,044 | 83 | 9 |
| ZINCpG000009hfT4 | Z6615016198 | 399,947 | 63,896 | 100 | 1 |
| ZINCov000006qjGm | Z6615179000 | 645,985 | 228,104 | 75 | 25 |
| ZINCoF000003N4Sx** | Z6615017403 | 880,786 | 626,545 | 103 | 25 |
| ZINCpK000006AHwY* | Z6615155291 | 936,860 | 370,458 | 71 | 6 |
| ZINClm00000kezXr | Z2775043007 | 2,629,858 | 2,412,686 | 97 | 18 |
| ZINCnA000000unIO | Z2027061106 | 502,284 | 20,893 | 97 | 18 |
| ZINCpC000007adah | Z6615015320 | 666,472 | 86,654 | 104 | 11 |
| ZINCr600000ge0zX | Z6615020155 | 946,329 | 248,727 | 128 | 22 |
| ZINCnI000005Jbfy | Z6615015266 | 13,220,440 | 8,857,701 | 99 | 45 |
| ZINCnr000000trh7 | Z6615020857 | 5,840,397 | 9,099,236 | 110 | 8 |
| ZINCpE0000069rny | Z6722169071 | 1,369,037 | 441,038 | 122 | 37 |
| ZINCoF000002XFGx | Z6615017320 | 591,813 | 109,443 | 97 | 41 |
| ZINCnx000002kQMC | Z2460724346 | 15,585,998 | 3,784,682 | 79 | 6 |
| ZINCns000000LyBi | Z2897416140 | 713,062 | 331,528 | 88 | 32 |
| ZINCnx000002sjNF | Z2610488449 | 1,285,283 | 158,771 | 95 | 25 |
| ZINCrA000007JHPl | Z6615018442 | 1,299,446 | 544,439 | 93 | 23 |
| ZINCoy0000006JE9 | Z2607647274 | 709,961 | 234,149 | 95 | 23 |
| ZINCot000004AncV | Z6615017518 | 1,749,614 | 618,223 | 100 | 17 |
| ZINCoA000004YwqJ | Z6717636067 | 559,191 | 104,829 | 93 | 10 |
| ZINCoz000005jTV2 | Z6615036658 | 25,027,141 | 3,099,950 | 135 | 16 |
| ZINCqo00000kNqT2 | Z6615022375 | 230,388,061 | 28,961,492 | 102 | 38 |
| ZINCpK0000061mQW | Z2940306468 | 848,656 | 487,238 | 88 | 4 |
| ZINCnN00000dISgx | Z6615018018 | 1,378,423 | 177,838 | 107 | 40 |
| ZINCps000005trlt | Z6615017520 | 1,409,088 | 588,227 | 96 | 7 |
| ZINCnN00000dIJkh | Z6615018562 | 590,593 | 102,350 | 89 | 52 |
| ZINCpO00000deSUE** | Z2940602274 | 408,463,466 | 71,344,460 | 67 | 15 |
| ZINCox000000PBvM | Z4070225106 | 546,045 | 42,858 | 82 | 4 |
| ZINCpR00000bUc1b | Z6615020280 | 826,997 | 205,535 | 76 | 18 |
| ZINCnA000000y1xj | Z3510328710 | 633,300 | 108,718 | 96 | 42 |
| ZINCov0000047szQ | Z6615016715 | 508,813 | 177,177 | 72 | 1 |
| ZINCot000004Aibf | Z6615022459 | 1,381,993 | 1,064,205 | 93 | 22 |
| ZINCoN00000e95OH | Z6615020105 | 644,036 | 281,891 | 114 | 29 |
| ZINCoB0000041653 | Z6615017597 | 1,478,976 | 147,356 | 144 | 8 |
| ZINCoO00000eidon | Z6615020027 | 1,058,206 | 342,940 | 103 | 9 |
| ZINCoM00000e5aKW | Z6615016774 | 1,236,436 | 279,771 | 85 | 13 |
| ZINCor000000eyEb | Z6615017908 | 1,441,621 | 833,580 | 104 | 21 |
| ZINCnw00000an5Mi | Z6615022396 | 11,212,019 | 1,114,704 | 99 | 33 |
| ZINCpi00000dt3Eq | Z6615020109 | 3,138,099 | 1,359,441 | 95 | 23 |
| ZINCoA000007zBqb | Z6615022393 | 739,277 | 78,847 | 83 | 5 |
| ZINCou000005LBQ1 | Z4462773688 | 955,325 | 211,989 | 93 | 10 |
| ZINCpz000006Bchs | Z2596056833 | 1,655,100 | 1,905,247 | 81 | 8 |
| ZINCpv000006LkJx | Z6615020916 | 403,911 | 57,771 | 100 | 17 |
| ZINCo9000009eD1K | Z6615018801 | 121,665,025 | 5,404,553 | 94 | 18 |
| ZINCox000008HLjQ | Z6615019214 | 2,446,020 | 490,972 | 110 | 10 |
| ZINCnC000007hEMI | Z6615016008 | 670,544 | 392,268 | 108 | 9 |
| ZINCpg00000D5L8V | Z6615014748 | 430,142 | 171,380 | 104 | 23 |
| ZINCoN00000eb66T | Z6615014610 | 374,120 | 57,229 | 77 | 40 |
| ZINCpJ00000btuG9 | Z6615015564 | 3,353,692 | 1,243,853 | 127 | 6 |
| ZINCoE000003wOdW | Z6615016222 | 1,034,615 | 595,714 | 71 | 13 |
| ZINCpw000005S8oM | Z6615124381 | 8,877,606 | 4,478,338 | 93 | 8 |
| ZINCoA000000msx0 | Z5389130126 | 476,887 | 212,614 | 75 | 10 |
| ZINCpf00000fKBYM | Z7044112626 | 597,730 | 261,485 | 95 | 0 |
| ZINCno00000blLI2 | Z2938832002 | 2,145,575 | 141,291 | 92 | 15 |
| ZINCop00000kXEki | Z6615122102 | 1,575,646 | 221,771 | 76 | 6 |
| ZINCox000008O53Q | Z6615020040 | 902,689 | 458,739 | 125 | 18 |
| ZINCpz000006XWGo | Z2940308998 | 557,638 | 224,054 | 78 | 14 |
| ZINCrp00000ciZ9T | Z6615015829 | 373,253 | 79,370 | 98 | 35 |
| ZINCql00000kMNvJ | Z6615020263 | 751,000 | 120,346 | 75 | 15 |
| ZINCnw00000290xU | Z6615017593 | 283,996 | 101,540 | 123 | 7 |
| ZINCoB00000415oH | Z3035221028 | 317,170 | 85,864 | 86 | 12 |
| ZINCpg00000ChI0z | Z6615014657 | 390,985 | 122,153 | 101 | 14 |
| ZINCoy00000077SN | Z6615020449 | 324,665 | 37,322 | 91 | 4 |
| ZINCos000004A5jq | Z6615022373 | 296,141 | 30,470 | 105 | 27 |
| ZINCjx000003jTFo | Z2897747949 | 1,369,196 | 121,049 | 91 | 18 |
| ZINCrn00000ciy0l | Z6615015424 | 411,345 | 111,957 | 85 | 27 |
| ZINCpG000009xP5O | Z2234667624 | 463,564 | 97,621 | 92 | 16 |
| ZINCjw000005PC05 | Z6615014898 | 288,698,728 | 6,568,397 | 95 | 11 |
| ZINCnH000008bnOR | Z2275216423 | 1,026,154 | 73,970 | 83 | 19 |
| ZINCnH000008gL5D | Z6615017736 | 324,510 | 21,784 | 100 | 27 |
| ZINCoK000004aIKt | Z6615022618 | 337,745 | 89,982 | 74 | 11 |
| ZINCoB000003YFFK | Z6615018405 | 455,960 | 94,760 | 91 | 4 |
| ZINCpz000008YxG6 | Z6615022384 | 447,717 | 22,441 | 92 | 4 |
| ZINCpv000006JYBB | Z2437439797 | 557,671 | 275,787 | 88 | 15 |
| ZINCoK000006E8lc | Z6615022794 | 830,640 | 206,344 | 81 | 15 |
| ZINCoz000005mkio | Z2940313312 | 2,006,920 | 789,870 | 89 | 16 |
| ZINCpv0000009ARJ | Z6615022433 | 787,456 | 234,952 | 99 | 20 |
| ZINCnt0000097uQi | Z3621468212 | 388,393 | 124,233 | 122 | 45 |
| ZINCnI000005G2xf | Z2940321940 | 526,283 | 40,753 | 94 | 30 |
| ZINCjz000003xoS3 | Z1740346668 | 483,323 | 293,162 | 75 | 26 |
| ZINCou000005M8el | Z4151090571 | 896,217 | 126,573 | 102 | 37 |
| ZINCpK000006xUJw | Z2940315175 | 530,858 | 50,164 | 80 | 7 |
| ZINCoM00000cV88d | Z2940316600 | 397,110 | 75,549 | 74 | 14 |
| ZINCnD000007yffK | Z2275400787 | 267,011 | 64,057 | 81 | 15 |
| ZINCmn00000dzoy1 | Z6615014758 | 604,139 | 107,888 | 108 | 25 |
| ZINCky000004t6c3 | Z2774814916 | 620,517 | 296,017 | 179 | 84 |
| ZINCpB000006BIww | Z6615015094 | 411,482 | 158,242 | 71 | 15 |
| ZINCoF000003NqKt** | Z6615016404 | 1,065,608 | 758,223 | 73 | 18 |
| ZINCob000000Fu00 | Z6615021877 | 670,910 | 29,617 | 97 | 26 |
| ZINCoO00000cYJ2f | Z2940602639 | 411,163 | 34,085 | 148 | 16 |
| ZINCnO00000dMi90** | Z6615017509 | 768,675 | 207,902 | 110 | 24 |
| ZINClo00000ks46n | Z4556474285 | 996,448 | 466,104 | 112 | 2 |
| ZINCnA000006ySao | Z6615022372 | 532,055 | 74,481 | 84 | 26 |
| ZINCkv000004fXeT | Z6615015420 | 413,382 | 4,533 | 106 | 17 |
| ZINCox0000081DUA | Z6615017782 | 752,907 | 13,343 | 86 | 10 |
| ZINCnk00000lzzM3 | Z2773641003 | 589,276 | 91,147 | 84 | 10 |
| ZINCnr000007ZKYn | Z6615015331 | 634,412 | 68,320 | 120 | 54 |
| ZINCoE000003wOkf | Z6615019239 | 585,522 | 87,014 | 79 | 10 |
| ZINCq800000ckcrv | Z6615014498 | 555,524 | 145,830 | 93 | 10 |
| ZINCnm00000n526N | Z2774427474 | 325,632 | 11,452 | 97 | 12 |
| ZINCoz000007jdD8 | Z6615020558 | 404,326 | 46,999 | 76 | 23 |
| ZINCnA000006CqwN | Z6615018339 | 431,879 | 70,860 | 102 | 21 |
| ZINCnu000009DLzG | Z6615022422 | 2,795,324 | 286,259 | 71 | 18 |
| ZINCoL00000dWJgJ | Z6615020275 | 760,455 | 465,407 | 88 | 27 |
| ZINCpz000006Xcju | Z2940320529 | 549,928 | 69,410 | 83 | 19 |
| ZINCmG000004Z2EV | Z6615014692 | 628,303 | 319,953 | 81 | 30 |
| ZINCom0000097L68 | Z2506306476 | 461,591 | 127,029 | 83 | 9 |
| ZINCoF000003PSlq | Z6615022462 | 666,433 | 594,593 | 95 | 17 |
| ZINCnF000007YVNA | Z2027059604 | 701,495 | 251,862 | 81 | 23 |
| ZINCpO00000f9NDa | Z6615019508 | 417,198 | 204,236 | 81 | 13 |
| ZINCoF000003MXoh | Z2940600508 | 578,847 | 297,526 | 62 | 9 |
| ZINCoT00000bVbE1 | Z6615016069 | 616,576 | 185,065 | 72 | 22 |
| ZINCpI00000aW94w | Z6615020201 | 291,650 | 20,186 | 112 | 27 |
| ZINCpE0000069rqo** | Z6615017470 | 360,429 | 100,064 | 101 | 25 |
| ZINCjs000005H1Qr | Z3494387572 | 385,316 | 187,081 | 72 | 14 |
| ZINCnu000004H7pl | Z4000325340 | 444,668 | 66,512 | 95 | 18 |
| ZINCnE000005ursN | Z2940315182 | 337,704 | 42,643 | 73 | 24 |
| ZINCov000006QgaK | Z6615017459 | 318,803 | 105,942 | 90 | 11 |
| ZINCns000000Fwfj | Z2437463231 | 383,815 | 102,281 | 106 | 17 |
| ZINCpH000006mtN0 | Z6615019960 | 659,818 | 148,385 | 99 | 34 |
| ZINCsE000004DJtp | Z6615014427 | 228,780 | 18,184 | 95 | 8 |
| ZINCpE000000jqSd | Z6615017528 | 18,294,211 | 2,265,024 | 105 | 26 |
| ZINCpJ000006uXyi | Z2940321111 | 727,075 | 407,315 | 84 | 20 |
| ZINCsE000004DJtF | Z6615020170 | 538,656 | 187,229 | 94 | 24 |
| ZINCpu000003Z443 | Z2940318508 | 233,813 | 62,289 | 81 | 17 |
| ZINCqz0000079HfX | Z6615014622 | 198,965 | 4,643 | 81 | 16 |
| ZINCqw000005Dd2D | Z2940322517 | 405,142 | 59,990 | 78 | 19 |
| ZINCsE000004DJts | Z6615017739 | 498,094 | 134,044 | 89 | 36 |
| ZINCoB000003U6KG | Z6615015126 | 529,774 | 33,769 | 106 | 13 |
| ZINCol0000097Ab7 | Z6615014517 | 282,374 | 4,030 | 89 | 11 |
| ZINCnq000005Qj6N | Z6615018515 | 302,837 | 51,742 | 111 | 6 |
| ZINCny00000bwi3N | Z6615017493 | 207,648 | 15,251 | 86 | 17 |
| ZINCoC000009DrFk | Z6615017387 | 432,906 | 62,563 | 100 | 21 |

The aggregation assays were done with 50 mM KPi, pH 7.0 of each compound, 1% DMSO. The compounds are screened at 200 µM. All MDH activities were measured at 200 µM, 100 µM * or 40 µM **. The data are the mean ± SD from three technical replicates.

**Extended Data Table 3.** **Data collection and refinement statistics (molecular replacement)**

|  | Z4462773688 | Z6615017509 | Z8427841182 | Z6615020275 | Z6615017782 |
| --- | --- | --- | --- | --- | --- |
| **Data collection** |  |  |  |  |  |
| Space group | C 2 | C 2 | C 2 | C 2 | C 2 |
| Cell dimensions |  |  |  |  |  |
| *a*, *b*, *c* (Å) | 119.03, 78.17, 97.72 | 118.47, 77.2, 97.64 | 119.77, 77.03, 100.08 | 119.19, 76.28, 98.1 | 118.23 76.9 97.69 |
| α, β, γ (°) | 90.00, 115.48, 90.00 | 90.00, 116.16, 90.00 | 90.00, 115.47, 90.00 | 90.00, 116.00, 90.00 | 90 116.544 90 |
| Resolution (Å) | 58.93-1.7 (1.761-1.7) | 58.54-1.663 (1.722-1.663) | 58.87-2.9 (3.004-2.9) | 58.57-1.7  (1.761-1.7) | 58.45-1.55 (1.605-1.55) |
| *R*_merge_ | 0.02177(0.3899) | 0.01435 (0.08429) | 0.1716 (0.1993) | 0.01791 (0.09036) | 0.02917 (0.8336) |
| CC_1/2_ | 0.999 (0.772) | 1 (0.984) | 0.837 (0.828) | 0.999 (0.977) | 0.999 (0.439) |
| *I* / σ*I* | 17.92 (1.87) | 27.30 (7.36) | 2.88 (1.24) | 24.78 (7.04) | 14.17 (1.05) |
| Completeness (%) | 96.92 (75.73) | 97.95 (95.98) | 98.62 (89.15) | 91.69 (58.24) | 99.31 (95.75) |
| Redundancy | 1.99 (1.98) | 1.99 (1.87) | 1.98 (1.87) | 1.99 (1.99) | 1.99 (1.99) |
| **Refinement** |  |  |  |  |  |
| Resolution (Å) | 58.93-1.7 | 58.54-1.663 | 58.87-2.9 | 58.57-1.7 | 58.45-1.55 |
| No. reflections | 171781 | 182635 | 35902 | 158716 | 224627 |
| *R*_work_ / *R*_free_ | 0.1873/ 0.2130 | 0.1868/0.2100 | 0.3055/0.3747 | 0.1617/0.1876 | 0.1848/0.2163 |
| No. atoms |  |  |  |  |  |
| Protein | 5663 | 5513 | 5485 | 5688 | 5715 |
| Ligand/ion | 48 | 46 | 46 | 48 | 48 |
| Water | 538 | 483 | N/A | 831 | 608 |
| *B*-factors |  |  |  |  |  |
| Protein | 34.67 | 31.53 | 23.32 | 22.84 | 29.26 |
| Ligand/ion | 42.16 | 51.14 | 49.67 | 39.08 | 52.07 |
| Water | 40.69 | 38.20 | N/A | 33.99 | 38.56 |
| R.m.s. deviations |  |  |  |  |  |
| Bond lengths (Å) | 0.43 | 0.38 | 0.53 | 0.37 | 0.48 |
| Bond angles (°) | 0.59 | 0.57 | 0.72 | 0.60 | 0.65 |
| Ramachandran plot (%) |  |  |  |  |  |
| Favored | 98.59 | 98.29 | 94.46 | 98.3 | 98.31 |
| Allowed | 1.41 | 1.71 | 3.26 | 1.70 | 1.69 |
| Disallowed | 0.14 | 0 | 0.28 | 0 | 0 |
| PDB ID | 9C81 | 9C6P | 9C83 | 9C84 | 9C8J |

*One crystal was used for each structure. Values in parentheses are for highest-resolution shell.

**References:**

1	Alon, A. *et al.* Structures of the sigma(2) receptor enable docking for bioactive ligand discovery. *Nature* **600**, 759-764, doi:10.1038/s41586-021-04175-x (2021).

2	Lyu, J. *et al.* Ultra-large library docking for discovering new chemotypes. *Nature* **566**, 224-229, doi:10.1038/s41586-019-0917-9 (2019).
